# A Pandas complex adapted for piRNA-guided transposon silencing

**DOI:** 10.1101/608273

**Authors:** Kang Zhao, Sha Cheng, Na Miao, Ping Xu, Xiaohua Lu, Yuhan Zhang, Ming Wang, Xuan Ouyang, Xun Yuan, Weiwei Liu, Xin Lu, Peng Zhou, Jiaqi Gu, Yiqun Zhang, Ding Qiu, Zhaohui Jin, Chen Su, Chao Peng, Jian-Hua Wang, Meng-Qiu Dong, Youzhong Wan, Jinbiao Ma, Hong Cheng, Ying Huang, Yang Yu

**Author notes:** Contributed equally (Kang Zhao, Sha Cheng, Na Miao, Ping Xu, Xiaohua Lu, Yuhan Zhang).

## Abstract

The repression of transposons by the Piwi-interacting RNA (piRNA) pathway is essential to protect animal germ cells. In *Drosophila* ovaries, Panoramix (Panx) enforces transcriptional silencing by binding to the target-engaged Piwi-piRNA complex, although the precise mechanisms by which this occurs remain elusive. Here, we show that Panx functions together with a germline specific paralogue of a nuclear export factor, dNxf2, and its cofactor dNxt1 (p15), as a ternary complex to suppress transposon expression. Structural and functional analyses demonstrate that dNxf2 binds Panx via its UBA domain, which plays an important role in transposon silencing. Unexpectedly, dNxf2 interacts directly with dNxf1 (TAP), a general nuclear export factor. As a result, dNxf2 prevents dNxf1 from binding to the FG repeats of the nuclear pore complex, a process required for proper RNA export. Transient tethering of dNxf2 to nascent transcripts leads to their nuclear retention. Therefore, we propose that dNxf2 may function as a Pandas (<u>Pa</u>noramix-d<u>N</u>xf2 <u>d</u>ependent T<u>A</u>P/p15 <u>s</u>ilencing) complex, which counteracts the canonical RNA exporting machinery and restricts transposons to the nuclear peripheries. Our findings may have broader implications for understanding how RNA metabolism modulates epigenetic gene silencing and heterochromatin formation.

---

Transposons are highly abundant in eukaryotes and make up nearly half of the human genome. To maintain eukaryotic genome integrity, nascent transcripts of transposons are often targeted by nuclear Argonaute proteins for transcriptional gene silencing (TGS)^1–5^. In animal gonads, the PIWI-clade Argonautes guided by piRNAs (PIWI-interacting RNA) are thought to recognize nascent transposon transcripts and direct sequence-specific heterochromatin formation^1–5^. As a critical cofactor of *Drosophila* nuclear Piwi, Panoramix (Panx, also known as Silencio) links the target-engaged Piwi-piRNA complex to the general silencing machinery^6,7^. Enforced tethering of Panx to nascent transcripts leads to cotranscriptional silencing and correlates with deposition of histone H3 lysine 9 trimethylation (H3K9me3) marks^6,7^. However, the mechanism by which Panx mediates this repression remains unknown. An equally important question is why H3K9me3 marks are not always sufficient for transposon silencing^8^.

To understand this fundamental question, we cross-examined proteins that co-immunoprecipitated with Panx (Extended Data Fig. 1a) and piRNA pathway candidate genes from RNA interference (RNAi) screens^9–12^. Unexpectedly, dNxf2 was identified as a potential cofactor of Panx (Extended Data Fig. 1a-c). dNxf2 belongs to an evolutionarily conserved NXF (<u>n</u>uclear <u>e</u>xport <u>f</u>actor) family of proteins, yet depletion of dNxf2 has no effect on polyadenylated mRNA export^13,14^. Instead, dNxf2 and its cofactor dNxt1 (also known as p15) have both been identified in two published RNAi screens as being essential for transposon silencing^9,10^. Like Panx, dNxf2 is specifically expressed in female gonads (Extended Data Fig. 1d).

To validate the interaction between Panx and dNxf2, GFP immunoprecipitation was performed from the ovaries expressing GFP-Panx fusion proteins under its native promoter. Results of mass spectrometry (Extended Data Fig. 1b) and western blot analysis demonstrated that endogenous dNxf2 was associated with Panx (Fig. 1a). Likewise, dNxf2-Halo with a Halo-tag integrated into the C-terminus was able to precipitate endogenous Panx from Ovarian Somatic Cell (OSC) lysates (Fig. 1b and Extended Data Fig. 6e). Next, we tested whether dNxf2 is functionally required for Panx-mediated silencing. The luciferase transcripts with BoxB sites in their 3’ untranslated regions are repressed if λN-Panx is tethered^6,7^. The expression level of luciferase was measured upon germline-specific knockdowns of either dNxf2 or dNxt1 (Fig. 1c). Despite constitutive tethering of λN-Panx, loss of either dNxf2 or dNxt1 significantly weakens the ability of Panx to repress the reporter, as compared to the controls (Zuc or attp2, Fig. 1c). Consistent with the reporter derepression, transposon transcripts are elevated upon dNxf2 RNAi (Extended Data Fig. 1e). Taken together, our data suggests that dNxf2 and dNxt1 may function as a heterodimer, either with or downstream of Panx, to suppress transposon expression.

**Figure 1.**
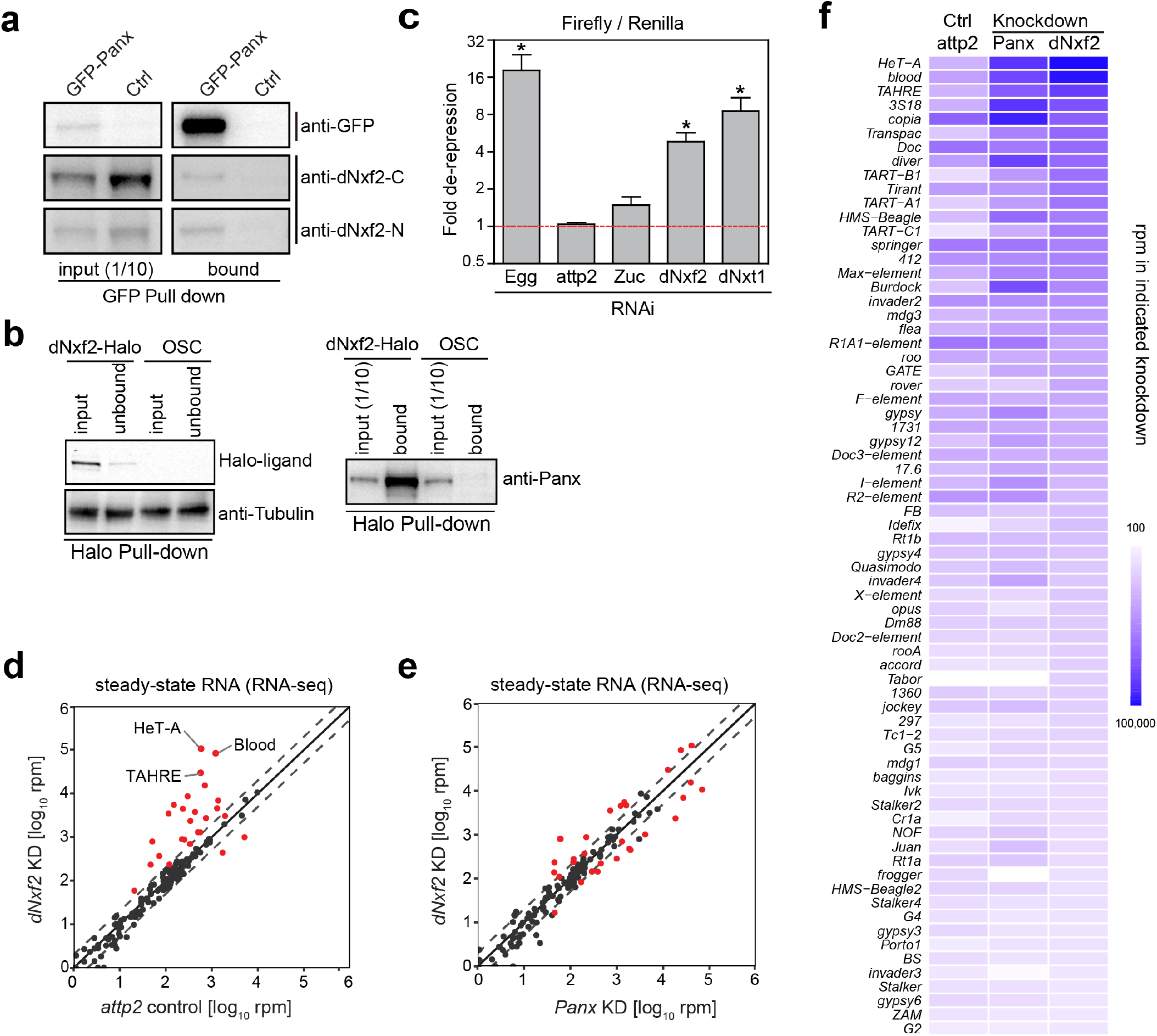
dNxf2 functions as a cofactor of Panx in the piRNA pathway. **a**, Western blots showing co-immunoprecipitation of endogenous dNxf2 with GFP-Panx from ovary lysates. Two different rabbit polyclonal monospecific dNxf2 (dNxf2-N and dNxf2-C) antibodies were used to detect endogenous dNxf2. **b**, Halo-ligand staining and western blots showing co-immunoprecipitation of endogenous Panx with dNxf2-Halo driven by the endogenous dNxf2 promoter from OSC lysates. A rabbit polyclonal monospecific Panx antibody was used to detect endogenous Panx. The left panel shows depletion of dNxf2-Halo proteins in the unbound sample by Halo beads, and the anti-Tubulin blots serve as loading controls; the right panel shows endogenous Panx proteins. **c**, The effects of germline (*nanos-*GAL4) knockdown of the indicated genes on Renilla-normalized Firefly luciferase activity of the reporter while tethering λN-Panx. For comparison, the relative value of the attp2 control was used for normalization. Data is shown as the mean ± s.d. (*n* = 15; \**p* = 1.41387E-07). **d**, Comparison of steady-state RNA levels are shown as reads per million (rpm) mapping to the sense strand of each transposon consensus from the *nanos*-GAL4 driven knockdown for dNxf2 (*Y* axis) versus control (*X* axis). Dashed lines indicate two-fold changes. The average of two replicates is shown. KD = knockdown. Red dots indicate transposon elements with significant changes. **e**, Comparison of steady-state RNA levels (RNA-seq; shown as RPM) mapping to the sense strands of each transposon consensus from the *nanos*-GAL4 driven knockdowns of the indicated genes. Red dots indicate transposon elements with significant changes from **d**. **f**, Heat map displaying steady-state RNA levels (RNA-seq) as reads per million (rpm) for the top 70 detected transposons from the *nanos*-GAL4 driven knockdowns of the indicated genes in a blue-white scale.

Next, we used RNA sequencing (RNA-seq) to examine global effects on transposon expression with germline-specific knockdowns of dNxf2, compared with Panx RNAi (Fig. 1d-f). As expected, dNxf2 knockdown triggered a dramatic increase of transposon transcripts (Fig. 1d), similar to that of Panx (Fig. 1e-f), suggesting that dNxf2 is specifically required for silencing of transposons repressed by Panx. To rule out off-target effects of RNAi, a loss of function mutant of dNxf2 was generated using CRISPR/Cas9 (Extended Data Fig. 2a)^15^. The dNxf2 mutant female flies carrying a deletion of 20 amino acids at the N-terminus were completely sterile (Extended Data Fig. 2b), similar to other core piRNA pathway mutants^6^. Yet, loss of dNxf2 showed little effect on Piwi nuclear localization or stability (Extended Data Fig. 2c-d), indicating that dNxf2 functions as an effector protein rather than in piRNA biogenesis. Consistent with this notion, the dNxf2 mutants showed global upregulation of transposons (Extended Data Fig. 2e-g) and derepression of the luciferase reporter, despite the λN-Panx tethering (Extended Data Fig. 3a). Unexpectively, absence of dNxf2 noticeably reduces endogenous Panx protein levels (Extended Data Fig. 2d). To rule out the possibility that dNxf2 may indirectly affect transposons via Panx stability, transposon expression levels were measured upon overexpression of λN-Flag-Panx under the dNxf2 mutant background (Extended Data Fig. 3a-e). Still, the dNxf2 mutant female flies lost transposon controls (Extended Data Fig. 3c-d) and were completely sterile, as if Panx did not exist (Extended Data Fig. 3e).

The striking phenotypic similarities between dNxf2 and Panx prompted us to test whether these two proteins interact directly. We used yeast two-hybrid (Y2H) assays to determine the interacting regions. The domain architecture of dNxf2 is very similar to that of the canonical RNA export factor, dNxf1 (also known as TAP or sbr, Fig. 2a). Both proteins contain leucine-rich repeats (LRR), an RNA recognition motif (RRM), a nuclear transport factor 2 (NTF2)-like domain, and a ubiquitin-associated domain (UBA). Interestingly, Panx only interacts with the UBA domain of dNxf2 but not that of dNxf1 (Fig. 2b), indicating that this interaction between Panx and dNxf2 is specific (Fig. 2b). Surprisingly, neither the full length nor the NTF2+UBA fragment of dNxf2 could bind Panx (Fig. 2b), suggesting that the UBA domain of dNxf2 might be in a “closed” conformation in the presence of the NTF2 domain. In this regard, the interactions between dNxt1 and the NTF2 domains of either dNxf2 or dNxf1 are also weakened in the presence of its UBA domain (Fig. 2c). Since *Drosophila* Nxt1 itself is absent in the Y2H system, we tested whether dNxt1 might release the UBA domain from the NTF2 domain to permit Panx binding. Indeed, ectopic expression of dNxt1 is sufficient to allow full length dNxf2 to interact with Panx in a Y2H assay (Fig. 2d). Next, we mapped the minimum region of Panx down to residues 315-343 (NIR, d<u>N</u>xf2 interacting <u>r</u>egion) as sufficient for UBA binding (Fig. 2e and Extended Data Fig. 3f). Consistent with the fact that dNxt1 forms a heterodimer with dNxf2^13^, we found that dNxt1 co-migrates with a fusion protein of dNxf2^NTF2+UBA^-(Gly-Ser)_4_-Panx^NIR^ by size-exclusion chromatography (Fig. 2f), suggesting that Panx, dNxf2, and dNxt1 may exist as a ternary complex. We were not able to crystalize the dNxf2^NTF2^ domain; instead, we crystallized dNxf1^NTF2^ in complex with dNxt1 and determined the structure (Extended Data Fig. 4a). Residues that may be involved in the binding of dNxf2^NTF2^ to dNxt1 were modeled according to the sequence alignment result and the structure of dNxf1^NTF2^ (Extended Data Fig. 4b). dNxf2^NTF2^ maintained most, if not all, residues that interact with dNxt1 as validated by the Y2H and co-immunoprecipitation assays (Fig. 2c and Extended Data Fig. 4a-e), indicating that the interaction mode of dNxf2^NTF2^/dNxt1 complex is like that of dNxf1^NTF2^/dNxt1.

**Figure 2.**
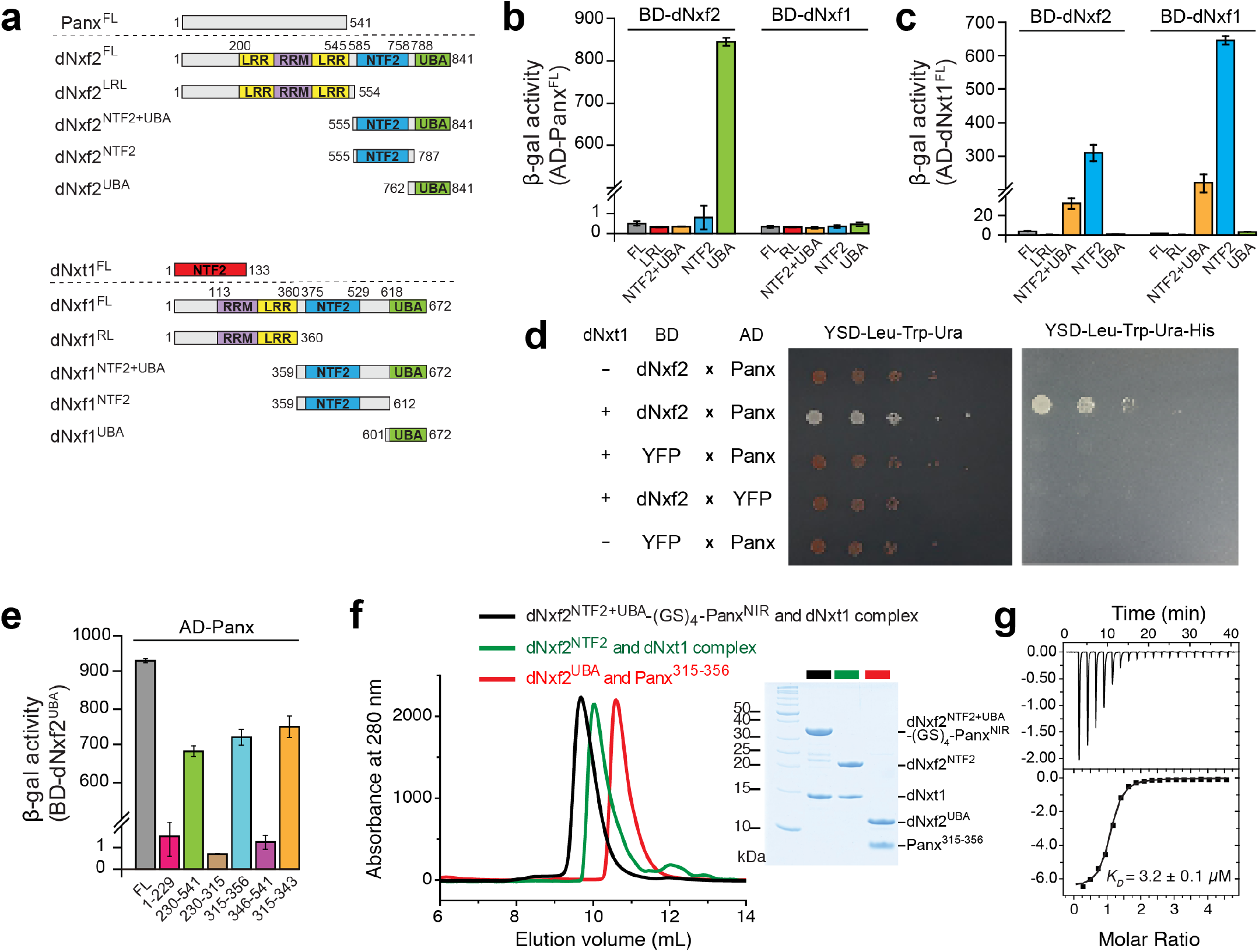
The UBA domain of dNxf2 interacts with Panx directly. **a**, Domain architectures of Panx, dNxf2, dNxf1, and dNxt1. Numbers above the diagrams correspond to amino acid residues of each protein. Domain names are abbreviated within respective colored regions. **b-e**, Yeast two-hybrid (Y2H) assays mapping the interacting regions between *Drosophila* Nxf1/2 and Panx or dNxt1. Interactions were determined by either measuring the beta-galactosidase activity produced by the reporter gene or growth on YSD media lacking the indicated essential amino acid or uracil. Data are averages of three independent experiments (n = 3). Proteins or fragments shown above the dashed line are used as preys in the assays. **b**, Y2H assays mapping regions of *Drosophila* Nxf1/2 that interact with Panx. **c**, Y2H assays mapping regions of *Drosophila* Nxf1/2 that interact with dNxt1. **d**, Yeast three hybrid assay determining the requirement of dNxt1 for a Nxf2:Panx interaction. **e**, Y2H assays mapping minimum regions of Panx that interact with dNxf2^UBA^. **f**, The left panel shows the size exclusion chromatography profile of the NTF2 and UBA domains of dNxf2 forming heterodimers with dNxt1 and Panx^NIR^ in solution, respectively. A dNxf2 fragment spanning the NTF2 and UBA domains that is covalently linked to Panx^NIR^ forms a stable complex with dNxt1. The right panel shows the components of the peak in the elution profile by SDS-PAGE. Color schemes used for the three complexes are indicated in the key. **g**, Quantification of the dissociation constant for the interaction between dNxf2^UBA^ and Panx^NIR^ as measured by an isothermal titration calorimetry assay.

Purified Panx^NIR^ forms a stable complex with the UBA domain of dNxf2 with a dissociation constant of ~3.2 μM as measured by isothermal titration calorimetry (ITC) (Fig. 2g). To further explore the molecular basis of interactions between dNxf2 and Panx, we determined the crystal structure of the dNxf2-Panx complex (Fig. 3a-d). Despite many efforts, only the fusion protein of dNxf2^UBA^-(Gly-Ser)_4_-Panx^NIR^ could successfully be crystallized. The structure was solved at 1.5 Å resolution (Extended Data Table 1). dNxf2^UBA^ forms a compact three-helix bundle (α1–α3) with a 3_10_-helix (η1) at the C-terminus (Fig. 3b). The Panx^NIR^ is folded into a long α-helix and lays on the hydrophobic surface formed by α2 and α3 (Fig. 3b-c and Extended Data Fig. 4f). A324, A328, V331, L332, and I335 on Panx interact with V800, F819, F826, F840, L823 and I827 on dNxf2 via hydrophobic interactions (Fig. 3d). Moreover, R321 and R327 on Panx form salt bridges with D837 and E799 on Nxf2^UBA^, respectively (Fig. 3d). To validate the intermolecular interactions between dNxf2 and Panx, key residues on the interacting interface were mutated (Fig. 3e). While either the L823A or D837A single point mutation affected the binding between Panx and dNxf2^UBA^, the double point mutation of dNxf2^UBA^ (F826A/I827A) nearly abolished its interaction with Panx in both Y2H and co-immunoprecipitation assays (Fig. 3e-f), highlighting the significant contribution of these residues in Panx binding.

**Figure 3.**
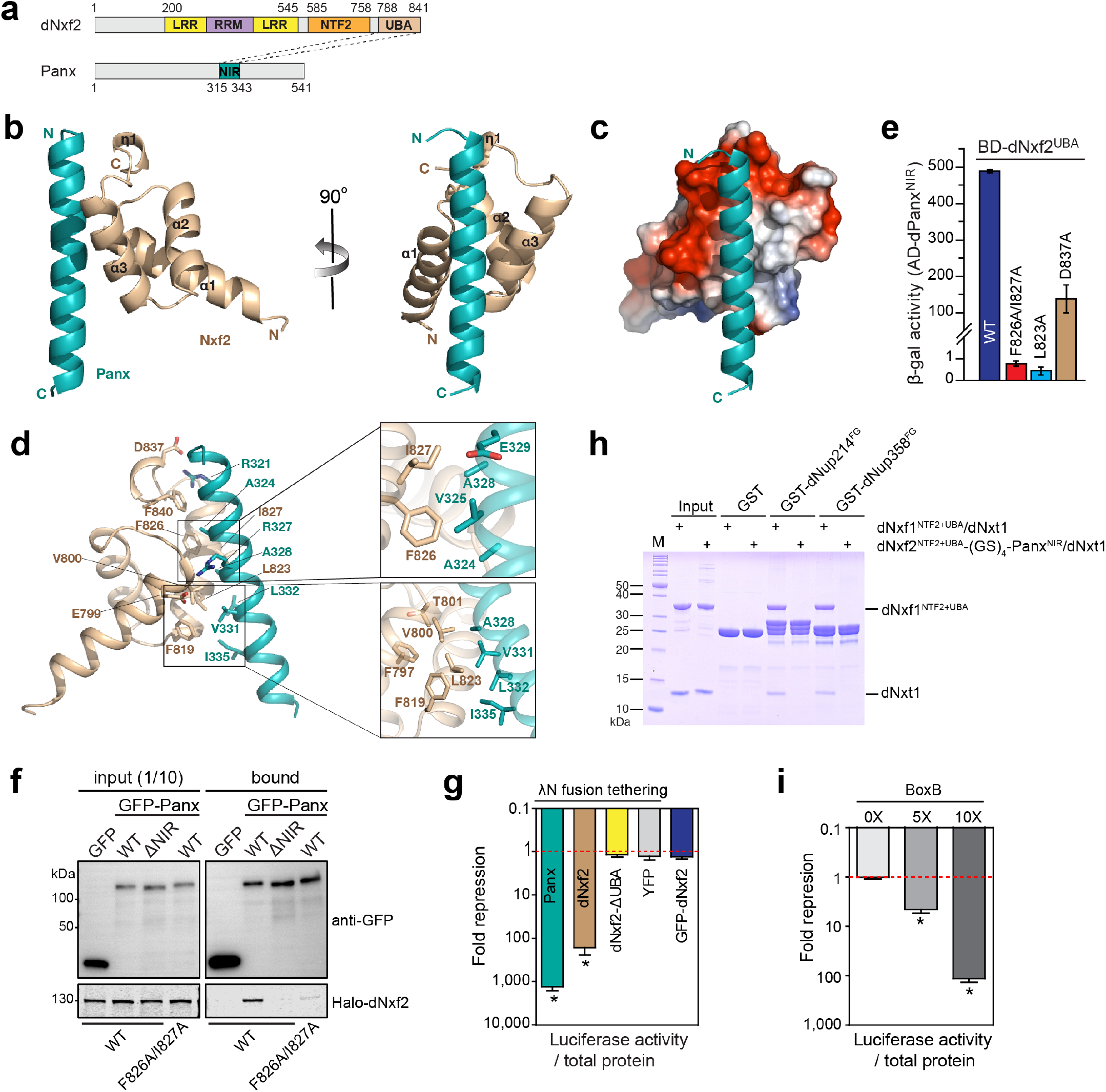
Structure of dNxf2^UBA^ in complex with Panx^NIR^. **a**, Schematic of the interacting region between dNxf2 and Panx. Numbers above or below the diagrams correspond to amino acid residues of dNxf2 or Panx, respectively. Domain names are abbreviated within respective colored regions. NIR, d<u>N</u>xf2 <u>i</u>nteracting region. **b**, Left, cartoon representation of dNxf2^UBA^ in complex with Panx^NIR^. The dNxf2^UBA^ and Panx^NIR^ fragment are colored in tan and teal, respectively. Right, a view rotated 90° around the vertical axis. **c**, Electrostatic potential analysis of the Panx-binding surface of dNxf2^UBA^. Panx^NIR^ is shown in cartoon mode. **d**, A detailed view of the interactions between dNxf2^UBA^ and Panx^NIR^. Key residues involved in binding are shown in sticks. Close-up views of hydrophobic interactions between dNxf2^UBA^ and Panx^NIR^ are shown on the right. **e**, Y2H assays measuring the binding of wild-type or mutant dNxf2^UBA^ with Panx^NIR^. Mutations of key residues are indicated along the bars. **f**, Western blots and Halo-ligand staining showing co-immunoprecipitation of GFP-tagged Panx or its NIR deletion mutant (ΔNIR) with Halo-tagged dNxf2 or its F826A/I827A double mutant from OSC cells. GFP serves as a negative control. **g**, Effects of the indicated λN fusion proteins or a non-tethering control (GFP-dNxf2) on luciferase activity of the reporters integrated into the attP2 landing site. Data is shown as the mean ± s.d. (*n* = 15; \**p* = 1.41387E-07). **h**, SDS-PAGE showing pull-down results of the dNxf1^NTF2+UBA^/dNxt1 complex and dNxf2^NTF2+UBA^-(GS)_4_-Panx^NIR^/dNxt1 complex by either GST-tagged dNup214^FG^ or dNup358^FG^, respectively, compared to a GST control. The beads were washed and aliquots of the bound fraction (20%) were analyzed. 2 μg of each input protein was loaded. Positions of molecular weight markers are indicated on the left in kDa. **i**, The effects of λN-dNxf2 tethering on luciferase activity of reporters with increasing number of BoxB sites. All reporters are integrated into the same genomic locus (attP2 landing site). Fold repression is calculated as total protein-normalized Firefly luciferase luminescent values of the control (no tethering) divided by that of the indicated experiments. Data is shown as the mean ± s.d. (*n* = 15; \**p* = 1.41387E-07).

In contrast to the highly charged surface of the Nxf1-type UBA (for example human Nxf1/hsNxf1 or yeast Mex67/scMex67), dNxf2^UBA^ favors hydrophobic binding with Panx (Fig. 3c and Extended Data Fig. 5a-b). Key residues on the interacting interface are highly conserved among different *Drosophila* species but altered in the Nxf1-type UBA (Extended Data Fig. 5a). On the opposite surface of the Nxf1-type UBA, a hydrophobic pocket is formed to accommodate the FxFG peptide of the nuclear pore complex (NPC) (Extended Data Fig. 5c-d). However, this pocket is missing in dNxf2^UBA^ due to a salt bridge formed between K829 and E814 (Extended Data Fig. 5c). Additionally, the bulky side chain of L825 on dNxf2^UBA^ may hinder FG binding (Extended Data Fig. 5c). In contrast, the corresponding amino acids in hsNxf1 (A602) or scMex67 (G583) contained much smaller side chains (Extended Data Fig. 5c-d), therefore, giving space for FxFG interactions. Consistent with the structural predictions, the dNxf2^UBA^ domain was unable to bind to the FG-repeats of dNup214, a NPC component known to interact with dNxf1^UBA^ in both Y2H and GST pull down assays (Fig. 3h and Extended Data Fig. 5e)^14^. Similar GST pull down results were obtained using the FG-repeats from dNup358 (Fig. 3h). Since two copies of FG binding domains (NTF2 and UBA) are minimally required for proper RNA export^16^, dNxf2 lacks at least one copy of the FG binding domain (UBA) and thus cannot export RNAs.

To validate the importance of the direct interactions between dNxf2 and Panx *in vivo*, a previously described λN/BoxB luciferase reporter system^6^ was used to check if artificial tethering of dNxf2 could lead to repression. As expected, significant repression upon tethering of a λN-dNxf2 fusion protein was observed (Fig. 3g), unlike that of the negative controls (λN-YFP or GFP-dNxf2 lacking a λN-tag). Like Panx, the level of λN-dNxf2 mediated repression was found in a dosage-dependent manner, which is correlated with the number of BoxB binding sites (Fig. 3i). Most importantly, the repression is dependent on the presence of the dNxf2 UBA domain (Fig. 3g, dNxf2-∆UBA).

Like dNxf1, dNxf2 contains RNA binding domains (RBDs) at the N-terminus, implying that dNxf2 might directly bind to transposon transcripts (Fig. 3a)^13^. To test this hypothesis, we performed GoldCLIP (Gel-ommitted Ligation-dependent CLIP)/RT-qPCR experiments, which rely on the covalent attachment of a Halo-tag to its ligand on beads to allow denaturing purification of crosslinked protein-RNA complexes^17^. A Halo-tag was inserted at the C-terminus of dNxf2 (dNxf2-Halo) using CRISPR/Cas9 (Extended Data Fig. 6a-f)^18^. Mdg1 is one of the transposon families targeted by Piwi/piRNAs in OSCs^8,19^. Strikingly, after UV crosslinking and denaturing washes, the transcripts of mdg1, but not the housekeeping gene rp49, remained attached to the dNxf2-Halo fusion protein (Extended Data Fig. 6b-c). This association depended on UV crosslinking, demonstrating a direct binding between mdg1 and dNxf2 (Extended Data Fig. 6b-c). Interestingly, the interactions were only observed when both histone H1 and heterochromatin protein 1a (HP1a) were depleted by RNAi, but not in the control knockdown (Extended Data Fig. 6b-c), supporting the idea that the majority of transposon transcripts remain suppressed in a wildtype background. Low steady-state levels of transposon nascent RNAs make it more difficult to obtain significant signals in an already inefficient UV crosslinking experiment. Nevertheless, upon removal of the downstream silencing factors (H1 and HP1a), transposon transcripts accumulate and are bound by dNxf2-Halo (Extended Data Fig. 6b). In contrast, Frogger, a transposon known not to be targeted by piRNAs^19^, did not show any detectable CLIP signal, although its transcripts are dramatically upregulated upon the H1/HP1a double knockdowns (Extended Data Fig. 6d). This result suggests that the binding of dNxf2 to transposons could be correlated with piRNA targeting^19^. Furthermore, we performed GoldCLIP-seq experiments using dNxf2-Halo knock-in OSCs depleted of Maelstrom, a piRNA pathway effector component either parallel or downstream of H3K9me3 establishment on transposons (personal communication with Mikiko Siomi)^8,20^. Consistent with the RT-qPCR results, CLIP-seq data supported the idea that dNxf2 preferentially binds to Piwi-targeted transposons, especially in the absence of Mael (Extended Data Fig. 7). Moreover, the RBDs of dNxf2 are essential for silencing in the tethering assays (Extended Data Fig. 8). Therefore, dNxf2^RBD^ is likely to be involved in the effector step of silencing rather than the Piwi-dependent recruitment of dNxf2. Collectively, our data is consistent with the model that Panx and dNxf2/dNxt1 function together as a stable complex to directly suppress transposons that are targeted by Piwi-piRNAs.

Since loss of Panx leads to a significant reduction of H3K9me3 marks on transposons^6,7^, we next tested whether the removal of dNxf2 could result in a similar phenomenon. Considering that Panx is unstable in the absence of dNxf2 (Extended Data Fig. 2d), we performed H3K9me3 ChIP-qPCR assays over several transposons as well as the Firefly-10xBoxB reporter, while overexpressing λN-Flag-Panx. H3K9me3 showed marginal changes upon removal of dNxf2. In contrast, transposon transcripts are still dramatically upregulated in the absence of dNxf2 (Extended Data Fig. 3c-d and 9). This result suggested that transposon silencing and H3K9me3 deposition could somehow be uncoupled in the dNxf2 mutant.

Interestingly, we found that HeT-A chromatin left nuclear peripheries upon loss of either Panx or dNxf2 (Extended Data Fig. 10)^21^. In this regard, I element transcripts, which are targeted by piRNAs, have previously been shown to accumulate within the nucleus^22^. Thus, we proposed that certain RNA export machineries may be regulated by the piRNA pathway to prevent transposon transcripts from being exported out of the nucleus. dNxf1 (TAP) could be one such candidate since hsNxf1 has been reported to dimerize with most NXF family members to regulate RNA export^23,24^. Therefore, we tested whether dNxf2 might interact with dNxf1 and counteract the RNA exporting function of dNxf1. Indeed, GFP-tagged dNxf2 can co-immunoprecipitate Halo-tagged dNxf1 from OSC lysates (Fig. 4a). Either the NTF2 or the UBA domain of dNxf1 is sufficient to interact with dNxf2^NTF2^ (Fig. 4b and Extended Data Fig. 11a-c). We noticed here that full length dNxf2 interacts rather weakly with dNxf1 (Fig. 4a), while the interactions seemed much stronger using the truncated versions (Fig. 4b and Extended Data Fig. 11a-c). This implied that the majority of dNxf1 are not available for dNxf2 binding. Consistent with the co-immunoprecipitation results, the interaction between dNxf2 and dNxf1 is sufficient to bring together a split Gaussia luciferase, strongly arguing that dNxf2 and dNxf1 can be in close proximity *in vivo* (Fig. 4c). Furthermore, the GST pull-down assays demonstrated a direct binding between dNxf1 and dNxf2 (Fig. 4d and Extended Data Fig. 12a-d). Using chemical crosslinking of proteins coupled with mass spectrometry^25^, we were able to identify key residues in the intermolecular crosslinks within the dNxf1/dNxf2/dNxt1 complex (Fig. 4e). We noticed here that K640 and K664 from the UBA domain of dNxf1, sitting on each side of the FG binding pocket, were crosslinked with dNxf2^NTF2^ (Extended Data Fig. 12e). This raised the possibility that dNxf2 binding might block the entry of FG repeats to dNxf1^UBA^. In fact, an excess amount of dNxf2^NTF2^ can compete dNxf1^NTF2+UBA^ off from the FG repeats of dNup214 (Fig. 4f). Taken together, our results strongly suggest that the dNxf2^NTF2^ domain can directly block the access of the FG binding pockets on either NTF2 or UBA domain of dNxf1. As the binding of dNxf1 to NPC FG repeats is essential for RNA export^13^, dNxf2 may inhibit transposon export through blocking dNxf1’s ability to bind NPC.

**Figure 4.**
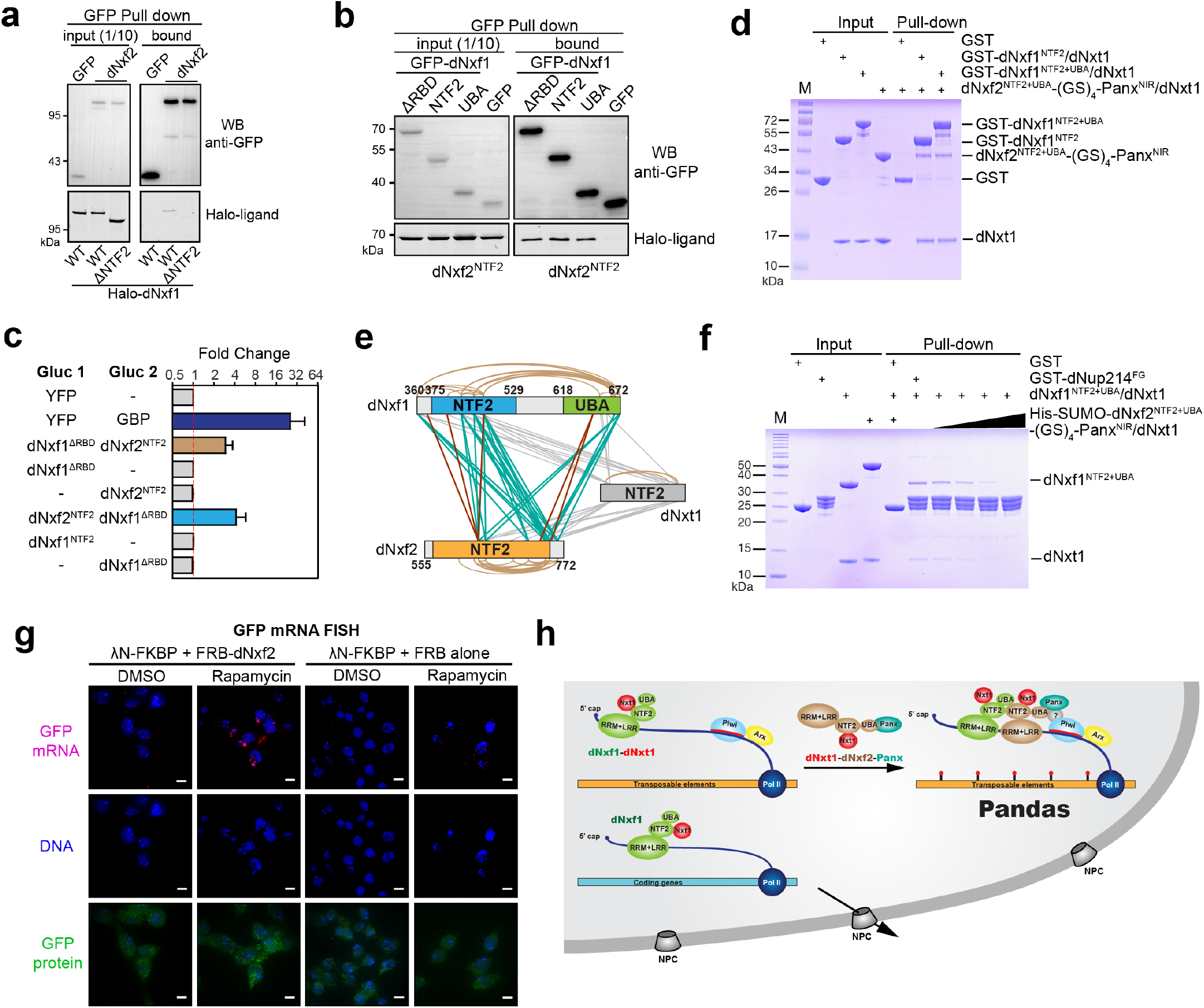
The Pandas complex is required for piRNA-guided transposon silencing. **a**, Western blots and Halo-ligand staining showing co-immunoprecipitation of GFP-tagged dNxf2 with Halo-tagged dNxf1 from OSC cells. ΔNTF2, Halo-dNxf1 lacking the NTF2 domain, and GFP serves as a negative control. **b**, Western blots and Halo-ligand staining showing co-immunoprecipitation of Halo-tagged dNxf2-NTF2 domain with different domain truncations of GFP-tagged dNxf1 from OSC cells. ΔRBD, dNxf1 lacking the N-terminus RRM and LRR domains; GFP serves as a negative control. **c**, Split Gaussia luciferase complementation assay results showing for coexpression of the indicated proteins in S2 cells. Fold changes are calculated as total protein normalized luciferase readings divided by that of the corresponding controls. Mean values ± s.d. from three independent experiments are shown. **d**, SDS-PAGE showing pull-down results of the Nxf2^NTF2+UBA^-(GS)_4_-Panx^NIR^/dNxt1 complex by either GST-tagged dNxf1^NTF2^/dNxt1 or dNxf1^NTF2+UBA^/dNxt1 respectively, compared to a GST control. The beads were washed and aliquots of the bound fraction (8%) were analyzed. Each input protein of 2 μg was loaded. Positions of molecular weight markers are indicated on the left in kDa. **e**, Schematic summary of statistical significant crosslinking residues identified between dNxf1^NTF2+UBA^, dNxf2^NTF2^ and dNxt1 in a recombinant complex reconstituted *in vitro*. Intermolecular crosslinks are shown as straight lines in either teal (dNxf1:dNxf2) or grey (dNxt1:dNxf1 or dNxt1:dNxf2) for DSS crosslinking, and in maroon (dNxf1:dNxf2) for EDC crosslinking. Intramolecular crosslinks were shown in brown curves. **f**, SDS-PAGE showing the competition assay results of the proteins precipitated by the GST-dNup214^FG^ pre-loaded with the dNxf1^NTF2+UBA^/dNxt1 complex, incubated with an increasing amount of the dNxf2^NTF2+UBA^-(GS)_4_-Panx^NIR^/dNxt1 complex. The beads were washed and aliquots of the bound fraction (20%) were analyzed. 2 μg of each input protein was loaded. Positions of molecular weight markers are indicated on the left in kDa. **g**, SIM super-resolution microscopy of RNA FISH. Comparison of the localizations of the reporter mRNAs tethered with λN-FKBP upon transient recruitment of either FRB-dNxf2 or FRB alone when treated with rapamycin for two hours to induce FKBP:FRB dimerization. DMSO serves as a control. Top panel, RNA signal (red) with DAPI staining; middle, DAPI staining (blue); bottom, RanGAP-GFP fusion protein (green). The scale bars represent 5 µm in length. **h**, A model for the Pandas (<u>Pa</u>noramix-d<u>N</u>xf2 <u>d</u>ependent T<u>A</u>P/p15 <u>s</u>ilencing) complex in piRNA-guided silencing, preventing nascent transposon RNA export when targeted by Piwi:piRNAs.

Next, we sought to directly visualize potential transient changes in RNA localization caused by dNxf2, using a rapamycin-inducible tethering system (Fig. 4g and Extended Data Fig. 13). Like any coding transcript, GFP mRNAs containing 10x copies of BoxB binding sites are mostly localized in the cytoplasm despite a constitutive λN-FKBP tethering (Fig. 4g and Movie S5-12). Upon rapamycin treatment, λN-FKBP dimerizes with the FRB-dNxf2 fusion protein, allowing a transient association of dNxf2 to the GFP mRNAs. Intriguingly, GFP mRNAs start to accumulate at nuclear peripheries upon binding of FRB-dNxf2 (Fig. 4g). The effect is specific to dNxf2 since FRB alone fails to cause any change. Given the involvement of Panx, dNxf2, dNxf1 (TAP) and dNxt1 (p15) in transposon silencing, we named this multi-protein complex as Pandas (<u>Pa</u>nx-d<u>N</u>xf2 <u>d</u>ependent T<u>A</u>P/p15 <u>s</u>ilencing). Our data raised the possibility that deterring the function of dNxf1 in transposon RNA export may be a key event in piRNA-guided silencing (Fig. 4h). In the absence of dNxf2, Panx fails to efficiently suppress either transposons or the tethered reporters (Extended Data Fig. 3a-e). In this regard, λN-Flag-Panx was unable to stay bound to the derepressed transposon chromatin as measured by Flag ChIP-qPCR (Extended Data Fig. 13c). Similarly, the Flag-ChIP signals over the tethered reporter were also diminished even though λN-Flag-Panx was constitutively tethered to the RNAs. It is well-established that HP1a can induce heterochromatin formation if tethered via DNA^26–29^. In contrast, direct tethering of HP1a to nascent RNAs fails to do so^7^. Therefore, our data provides a mechanistic insight that sequestering nascent transposon transcripts within the nucleus might be important to fully establish heterochromatin and enforce silencing. By removing dNxf2, we might have uncovered an intermediate state of silencing during heterochromatin formation.

Like any coding mRNA, transposon transcripts would likewise be transported into the cytoplasm by the general RNA exporting machinery (dNxf1/dNxt1), if not restrained by Piwi-piRNAs^14,22^. In piRNA-guided TGS, dNxf2 may function together with Panx as a stable complex to counteract this process (Fig. 4h). Our structure provides mechanistic insights into why dNxf2^UBA^ prefers to bind the silencing factor Panx rather than the FG repeats of NPCs (Extended Data Fig. 5). Remarkably, we found that dNxf2 can compete with the ability of dNxf1 to bind NPCs (Fig. 4f), thereby preventing RNA export. As dNxf2 preferentially associates with the piRNA-targeted transcripts (Extended Data Fig. 6-7), only a subset of dNxf1 associated with transposons could be affected by dNxf2. Accordingly, part of the silencing function of dNxf2 may be locally hijacking the RNA export machinery and repurposing dNxf1 into a “dead-end” complex, hence trapping transposon transcripts within the nuclear peripheries (Fig. 4h and Extended Data Fig. 13). Interestingly, Dam-ID has shown that both Piwi and NPCs contact chromatin at similar regions^30^. In this regard, dNxf1 has been found to be localized to nuclear peripheries where most constitutive heterochromatin resides^13,31–33^. Our data suggested that sequestering transposons to nuclear peripheries via the Pandas complex may help to establish/maintain their heterochromatic state (Extended Data Fig. 10 and Movie 1-4)^32,33^. Intriguingly, Xist can relocate the silenced X-chromosome to nuclear rim during X chromosome inactivation in mammalian cells, indicating that similar principles may apply to facultative heterochromatin formation marked with H3K27me3^34^. Recent evidence has demonstrated that hsNxf1 is required for efficient elongation of RNA polymerase II^35^. It is tempting to speculate that the Pandas complex might also inhibit transcriptional elongation of transposons via neutralizing dNxf1. In summary, we have uncovered an unexpected role of transposon RNA export blockage required for TGS. Our results will have broader implications for understanding how RNA metabolism modulates epigenetic gene silencing and heterochromatin formation (Fig. 4h).

## ACKNOWLEDGEMENTS

We thank the National Facility for Protein Science in Shanghai, Zhangjiang Lab, and Shanghai Science Research Center for their instrumental support and technical assistance. We thank the staff from BL19U1 beamline at the Shanghai Synchrotron Radiation Facility (SSRF) for assistance during data collection. We thank Ms. Shuoguo Li from the Center for Biological Imaging (CBI), Institute of Biophysics, Chinese Academy of Science for her help with taking and analyzing SIM images. This work was supported in part by grants from the Ministry of Science and Technology of China (2017YFA0504200 to Y.Y.) and from the National Natural Science Foundation of China (91640102 and 31870741 to Y.H.; 91640105 and 31770875 to Y.Y.), and the National Postdoctoral Program for Innovative Talents (BX20190081 to Y.H.Z.).

## AUTHOR CONTRIBUTIONS

Y.Y. and Y.H. conceived the project and wrote the manuscript. K.Z. constructed the dNxf2-Halo knock-in OSCs and established the dNxf2 mutant. K.Z., P.X., W.W.L., X.H.L., and D.Q. performed co-immunoprecipitations, tethering assays, transgenic fly constructions, RNA-seq, RT-qPCR experiments. S.C. and X.Y. performed structural studies, beta-gal activity assays, and ITC experiments; S.C. and Y.H.Z. performed GST pull-down assays. K.Z., Z.J., P.Z., X.O., J.G., and P.X. performed cloning. S.C. and X.L. performed Y2H assays. K.Z. and X.O. performed FACS analysis. N.M. preformed FISH and RNAi experiments. M.W. and Y.Q.Z. performed bioinformatics analysis. C.S., C.P., J.H.W., and M.Q.D. performed mass spectrometry and analyzed the data. Y.W., J.M. and H.C. provided critical reagents and advice; All authors discussed the results and commented on the manuscript.

## COMPETING INTERESTS

The authors declare that there is no conflict of interests.

## MATERIALS & CORRESPONDENCE

All materials and correspondence requesta should be addressed to.

## Notes

#### Summary of Updates

We added new evidences to demonstrate that dNxf2 interacts with dNxf1. Most importantly, we showed that dNxf2 can compete dNxf1 off from the FG repeats of Nup, providing key evidence how dNxf2 blocks dNxf1 from exporting nascent transposon RNAs.

## REFERENCES

1. Ge, D. T. & Zamore, P. D. Small RNA-directed silencing: the fly finds its inner fission yeast? Curr. Biol. 23, R318–20 (2013).

2. Martienssen, R. & Moazed, D. RNAi and heterochromatin assembly. Cold Spring Harb Perspect Biol 7, a019323 (2015).

3. Czech, B. & Hannon, G. J. One Loop to Rule Them All: The Ping-Pong Cycle and piRNA-Guided Silencing. Trends Biochem. Sci. 0, (2016).

4. Ozata, D. M., Gainetdinov, I., Zoch, A., Carroll, D. X. N. O. X. & Zamore, P. D. PIWI-interacting RNAs: small RNAs with big functions. Nat. Rev. Genet. 1–20 (2018). doi:10.1038/s41576-018-0073-3

5. Gainetdinov, I., Colpan, C., Arif, A., Cecchini, K. & Zamore, P. D. A Single Mechanism of Biogenesis, Initiated and Directed by PIWI Proteins, Explains piRNA Production in Most Animals. Mol. Cell 71, 775–790.e5 (2018).

6. Yu, Y. et al. Panoramix enforces piRNA-dependent cotranscriptional silencing. Science 350, 339–342 (2015).

7. Sienski, G. et al. Silencio/CG9754 connects the Piwi-piRNA complex to the cellular heterochromatin machinery. Genes Dev. 29, 2258–2271 (2015).

8. Sienski, G., Dönertas, D. & Brennecke, J. Transcriptional silencing of transposons by Piwi and maelstrom and its impact on chromatin state and gene expression. Cell 151, 964–980 (2012).

9. Czech, B., Preall, J. B., McGinn, J. & Hannon, G. J. A transcriptome-wide RNAi screen in the Drosophila ovary reveals factors of the germline piRNA pathway. Mol. Cell 50, 749–761 (2013).

10. Muerdter, F. et al. A genome-wide RNAi screen draws a genetic framework for transposon control and primary piRNA biogenesis in Drosophila. Mol. Cell 50, 736–748 (2013).

11. Handler, D. et al. The genetic makeup of the Drosophila piRNA pathway. Mol. Cell 50, 762–777 (2013).

12. Guruharsha, K. G. et al. A protein complex network of Drosophila melanogaster. Cell 147, 690–703 (2011).

13. Herold, A., Klymenko, T. & Izaurralde, E. NXF1/p15 heterodimers are essential for mRNA nuclear export in Drosophila. RNA 7, 1768–1780 (2001).

14. Katahira, J. Nuclear export of messenger RNA. Genes (Basel) 6, 163–184 (2015).

15. Port, F., Chen, H.-M., Lee, T. & Bullock, S. L. Optimized CRISPR/Cas tools for efficient germline and somatic genome engineering in Drosophila. Proc. Natl. Acad. Sci. U.S.A. 111, E2967–76 (2014).

16. Braun, I. C., Herold, A., Rode, M. & Izaurralde, E. Nuclear export of mRNA by TAP/NXF1 requires two nucleoporin-binding sites but not p15. Mol. Cell. Biol. 22, 5405–5418 (2002).

17. Gu, J. et al. GoldCLIP: Gel-omitted Ligation-dependent CLIP. Genomics Proteomics Bioinformatics 16, 136–143 (2018).

18. Savic, D. et al. CETCh-seq: CRISPR epitope tagging ChIP-seq of DNA-binding proteins. Genome Res. 25, 1581–1589 (2015).

19. Iwasaki, Y. W. et al. Piwi Modulates Chromatin Accessibility by Regulating Multiple Factors Including Histone H1 to Repress Transposons. Mol. Cell 63, 408–419 (2016).

20. Chang, T. H. et al. Maelstrom Represses Canonical Polymerase II Transcription within Bi- directional piRNA Clusters in Drosophila melanogaster. Mol. Cell 73, 291–303.e6 (2019).

21. Radion, E. et al. Key role of piRNAs in telomeric chromatin maintenance and telomere nuclear positioning in Drosophila germline. Epigenetics Chromatin 11, 40 (2018).

22. Chambeyron, S. et al. piRNA-mediated nuclear accumulation of retrotransposon transcripts in the Drosophila female germline. Proc. Natl. Acad. Sci. U.S.A. 105, 14964–14969 (2008).

23. Matzat, L. H., Berberoglu, S. & Lévesque, L. Formation of a Tap/NXF1 homotypic complex is mediated through the amino-terminal domain of Tap and enhances interaction with nucleoporins. Mol. Biol. Cell 19, 327–338 (2008).

24. Aibara, S., Katahira, J., Valkov, E. & Stewart, M. The principal mRNA nuclear export factor NXF1:NXT1 forms a symmetric binding platform that facilitates export of retroviral CTE-RNA. Nucleic Acids Res. 43, 1883–1893 (2015).

25. Combe, C. W., Fischer, L. & Rappsilber, J. xiNET: cross-link network maps with residue resolution. Mol. Cell Proteomics 14, 1137–1147 (2015).

26. Danzer, J. R. & Wallrath, L. L. Mechanisms of HP1-mediated gene silencing in Drosophila. Development 131, 3571–3580 (2004).

27. Hines, K. A. et al. Domains of heterochromatin protein 1 required for Drosophila melanogaster heterochromatin spreading. Genetics 182, 967–977 (2009).

28. Li, Y., Danzer, J. R., Alvarez, P., Belmont, A. S. & Wallrath, L. L. Effects of tethering HP1 to euchromatic regions of the Drosophila genome. Development 130, 1817–1824 (2003).

29. Azzaz, A. M. et al. Human heterochromatin protein 1α promotes nucleosome associations that drive chromatin condensation. J. Biol. Chem. 289, 6850–6861 (2014).

30. Ilyin, A. A. et al. Piwi interacts with chromatin at nuclear pores and promiscuously binds nuclear transcripts in Drosophila ovarian somatic cells. Nucleic Acids Res. 45, 7666–7680 (2017).

31. Kerkow, D. E. et al. The structure of the NXF2/NXT1 heterodimeric complex reveals the combined specificity and versatility of the NTF2-like fold. J. Mol. Biol. 415, 649–665 (2012).

32. van Steensel, B. & Belmont, A. S. Lamina-Associated Domains: Links with Chromosome Architecture, Heterochromatin, and Gene Repression. Cell 169, 780–791 (2017).

33. Towbin, B. D., Meister, P. & Gasser, S. M. The nuclear envelope--a scaffold for silencing. Curr. Opin. Genet. Dev. 19, 180–186 (2009).

34. Chen, C.-K. et al. Xist recruits the X chromosome to the nuclear lamina to enable chromosome-wide silencing. Science 354, 468–472 (2016).

35. Chen, S. et al. The mRNA Export Receptor NXF1 Coordinates Transcriptional Dynamics, Alternative Polyadenylation, and mRNA Export. Mol. Cell 74, 118–131.e7 (2019).

