## Supplementary Information for "A Pandas complex adapted for piRNA-guided transposon silencing"

### Extended Data Figure 1

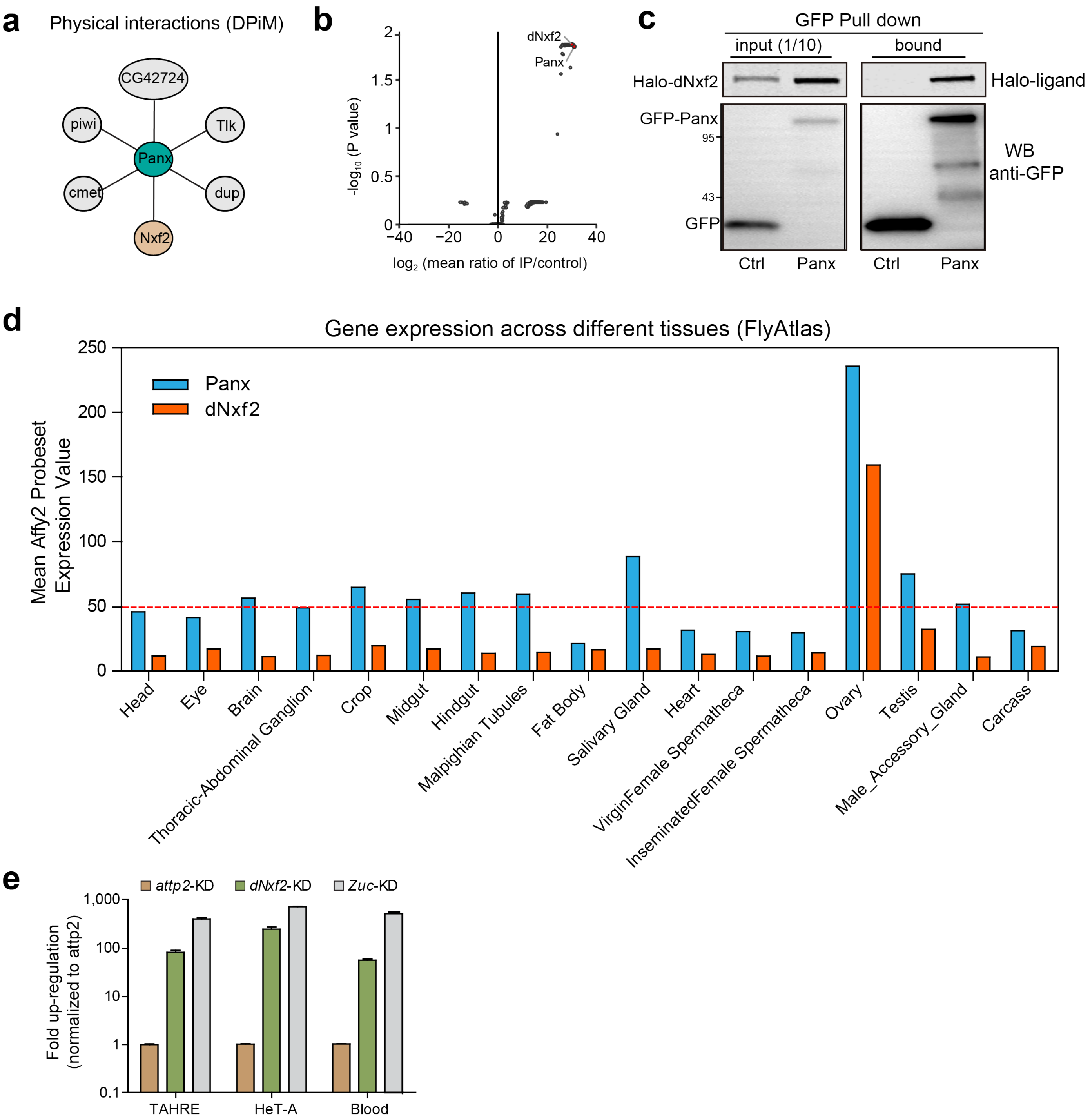


**Extended Data Figure 1. dNxf2 binds to Panx and is required for transposon silencing. a,** A network graph showing proteins co-immunoprecipitated with Flag-tagged Panx from the DPiM data downloaded from flybase.org. **b,** Volcano plots showing significance and enrichment values for proteins co-purified with GFP-Panx (n=2) from ovary lysates (the two most significantly enriched proteins are labeled). **c,** Western blots and Halo-ligand staining showing co-immunoprecipitation of GFP-tagged Panx with Halo-tagged dNxf2 from OSC cell lysates. GFP serves as a negative control. **d,** The expression profiles (*Y*-axis) of Panx and dNxf2 across different adult tissues (*X*-axis) are shown (blue, Panx; red, dNxf2). The mean affy2 probeset expression value (*X*-axis) is based on FlyAtlas anatomical microarray data downloaded from flybase.org. **e,** RT-qPCR results showing relative steady-state RNA levels of the indicated transposons for the germline specific (nanos-GAL4) knockdown of the indicated genes. The attp2 is used as a control**.** Fold changes are calculated as *rp49*-normalized RNA levels divided by that of the corresponding controls. Mean values ± s.d. from 3 independent experiments are shown.

### Extended Data Figure 2

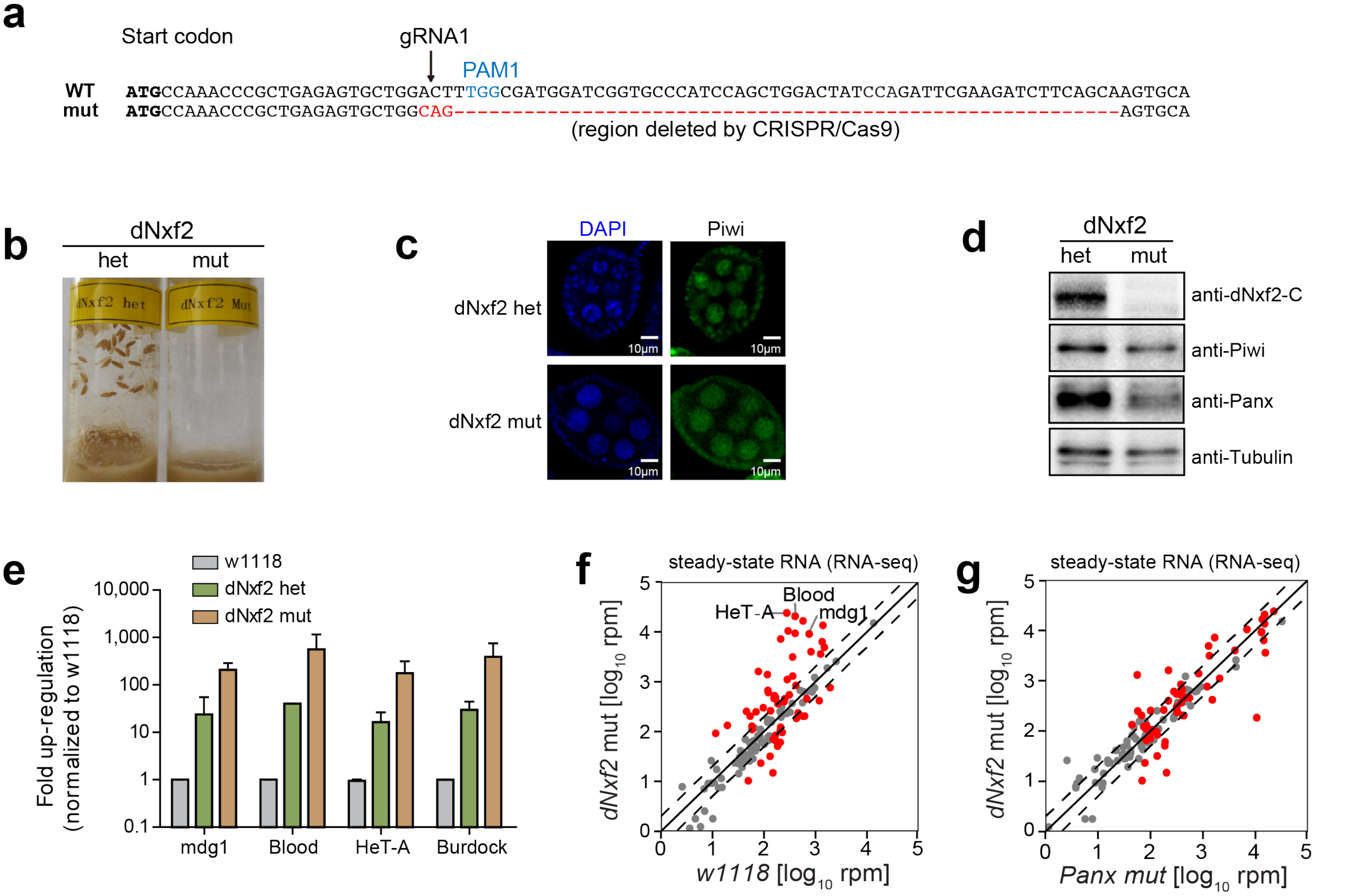


**Extended Data Figure 2. dNxf2 mutant resembles Panx mutant. a,** A schematic of CRISPR/Cas9-induced mutation in the open reading frame of dNxf2 is shown. Arrowheads indicate the cleavage sites targeted by the gRNA with the corresponding PAM site highlighted in light blue. The resulting mutant is shown below the wild-type sequence with the mutated/deleted region shown in red. **b,** Vials showing pupae from equal number of either heterozygous or mutant dNxf2 female flies crossed with OregonR males. **c,** Representative immunostaining of endogenous Piwi protein (green) from stage 5 ovaries of either heterozygous or mutant dNxf2. DAPI staining (blue) was used to indicate nuclei. **d,** Western blots showing protein expression level of endogenous dNxf2, Panx and Piwi from either heterozygous or mutant dNxf2 ovary lysates. Beta-tubulin serves as a loading control. **e,** RT-qPCR results showing relative steady-state RNA levels of the indicated transposons for the mutant or heterozygous dNxf2 ovaries, compared to the wildtype control (w1118). Fold changes are calculated as *rp49*-normalized RNA levels divided by that of the corresponding controls. Mean values ± s.d. from 3 independent experiments are shown. **f,** Comparison of steady-state RNA levels are shown as reads per million (rpm) mapping to the sense strands of each transposon consensus from dNxf2 mutant versus w1118 wild-type fly ovaries. Dashed lines indicate two-fold changes. The average of two replicates is shown. Red dots indicate transposon elements with significant changes. **g,** Comparison of steady-state RNA levels (RNA-seq; shown as RPM) mapping to the sense strands of each transposon consensus from dNxf2 mutant versus Panx mutant fly ovaries. Red dots indicate transposon elements with significant changes from **f**.

### Extended Data Figure 3

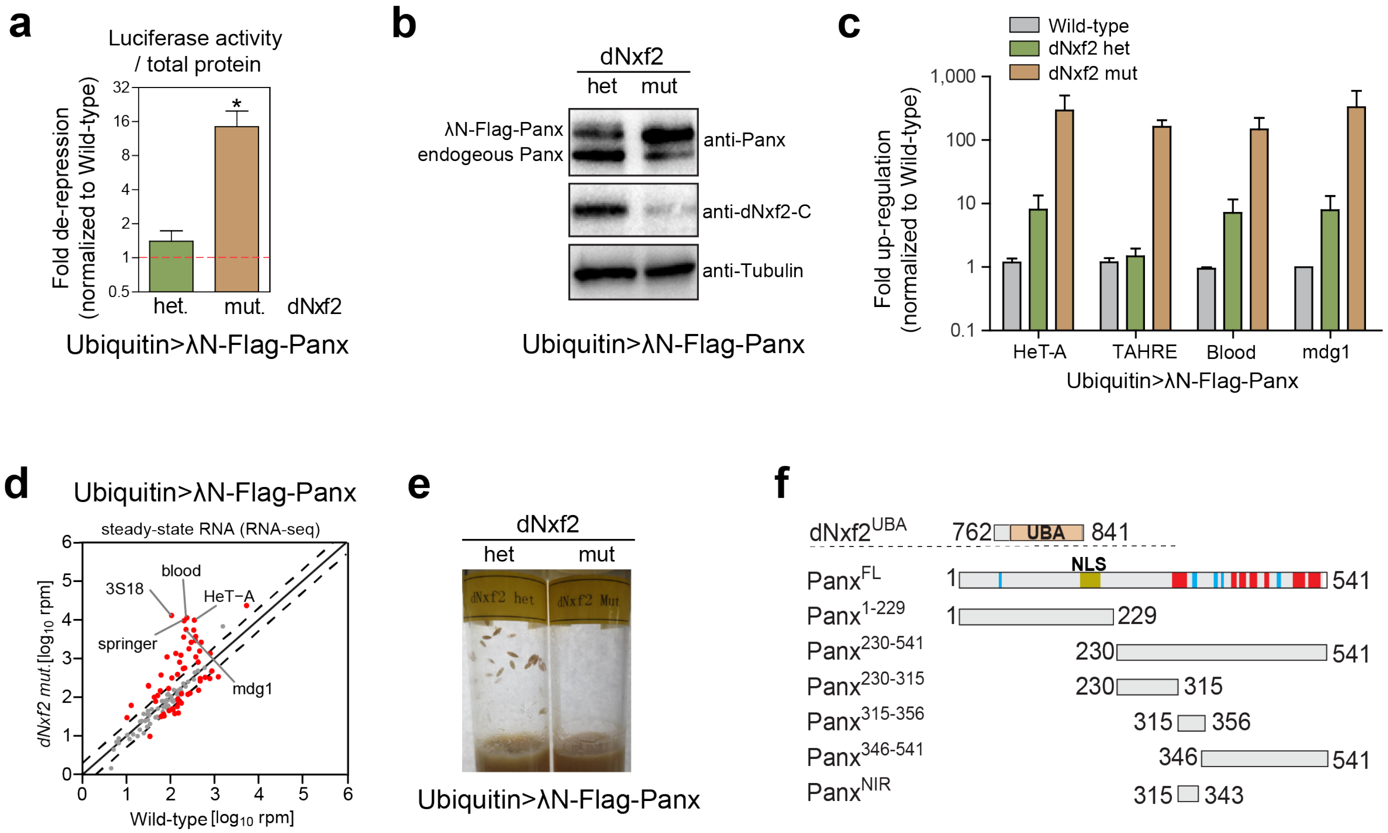


**Extended Data Figure 3. dNxf2 is required for Panx-mediated transposon silencing. a,** The expression of the total protein-normalized Firefly luciferase activity of the reporter under the indicated background (heterozygous control or transheterozygous mutant) while overexpressing λN-Flag-Panx driven by a Ubiquitin promoter. Data show mean ± s.d. ($n$ = 15; *$P$ = 1.41387E-07). **b,** Western blots showing protein expression level of endogenous dNxf2, Panx from either heterozygous or mutant dNxf2 ovary lysates while overexpressing λN-Flag-Panx driven by a Ubiquitin promoter. Beta-tubulin serves as a loading control. **c,** RT-qPCR results showing relative steady-state RNA levels of the indicated transposons for the mutant or heterozygous dNxf2 ovaries, compared to the wildtype control while overexpressing λN-Flag-Panx driven a Ubiquitin promoter. Fold changes are calculated as *rp49*-normalized RNA levels divided by that of the corresponding controls. Mean values ± s.d. from 3 independent experiments are shown. **d,** Comparison of steady-state RNA levels are shown as reads per million (rpm) mapping to the sense strands of each transposon consensus from dNxf2 mutant versus wild-type fly ovaries when overexpressing ubiquitin driven Panx. Dashed lines indicate two-fold changes. The average of two replicates is shown. Red dots indicate transposon elements with significant changes. **e,** Vials showing pupae from equal number of either heterozygous or mutant dNxf2 female flies when overexpressing ubiquitin driven Panx, mated with OregonR males. **f,** Schematic of full-length Panx and its truncated fragments used in the Y2H assays in Fig. 2e. Numbers corresponding to amino acid residues at either the N- or C-terminus of each protein fragment.

### Extended Data Figure 4

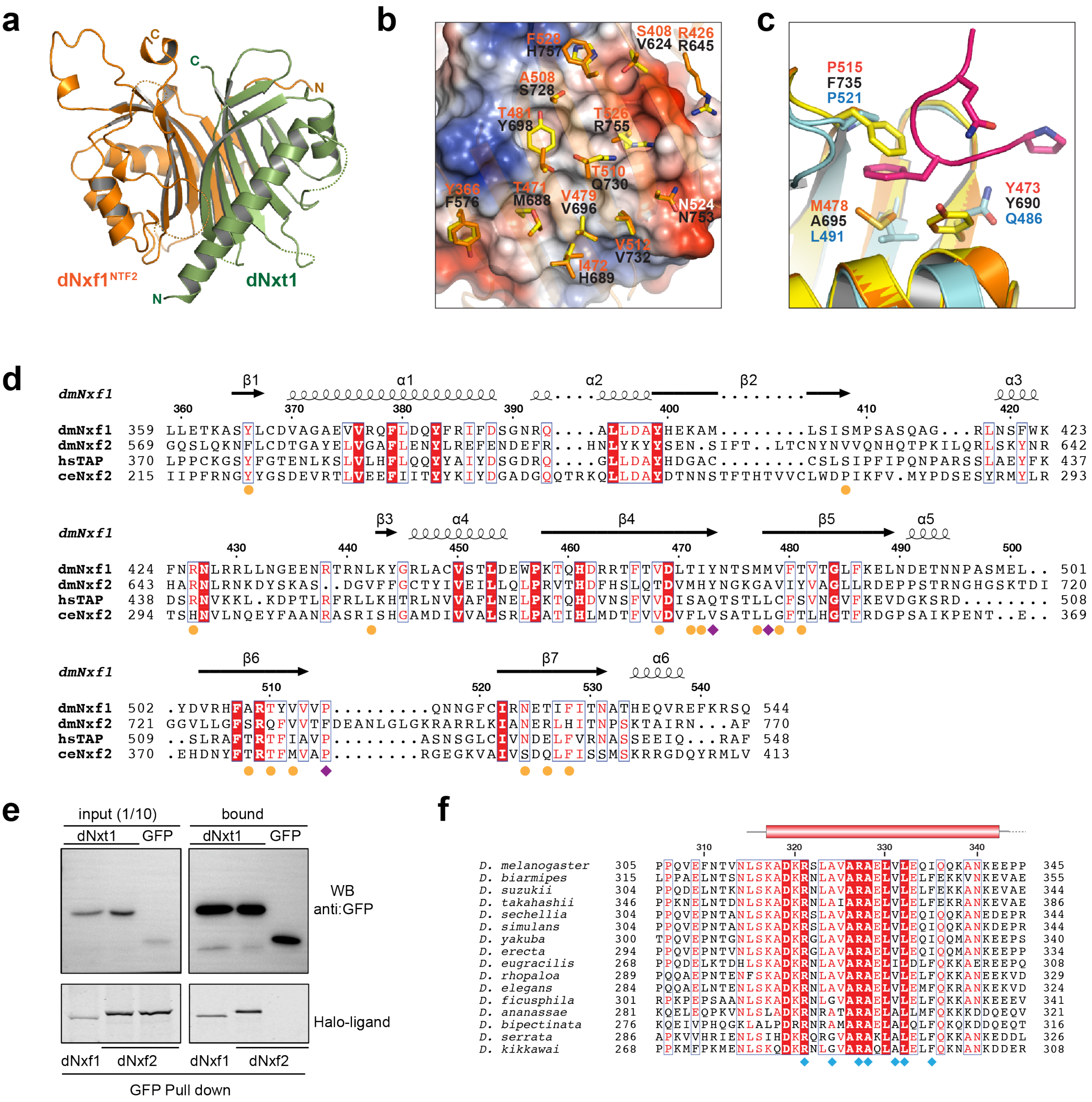


**Extended Data Figure 4. dNxf2 Structural analysis of dNxf1$^{NTF2}$ in complex with dNxt1. a,** Cartoon representation of dNxf1 NTF2 domain (orange) in complex with dNxt1 (green). **b,** Electrostatic surface potential representation of dNxt1, highlighting hydrophobic and electrostatic surfaces continuously from acidic (red) to non-polar (grey), to basic (blue). Key residues of dNxf1 are shown in stick mode (orange). The corresponding residues in dNxf2 (yellow) are modelled in Coot by mutation operation based on the dNxf1 structure in this study. **c,** dNxf1 NTF2 (orange) and dNxf2 NTF2 (yellow, model based on dNxf1 structure) are superimposed with human TAP NTF2 domain (cyan, PDB accession code 1JN5). Key residues that interact with the Phenylalanine-Glycine (FG) peptide (magenta) are labelled and shown in sticks. **d,** Sequence alignment of the NTF2 domains of dNxf1, dNxf2, human TAP (Q9UBU9), and ceNxf2 (Q9XVS8). The secondary structure annotation of dNxf1 is shown on the top. Residue positions of the reference sequence (dNxf1) are also indicated above the sequences. Arrows and spirals indicate beta-strands and alpha-helices, respectively. Red letters indicate similar residue; white letters in red shading indicate identical residues; blue boxes indicate conserved positions. **e,** Western blots and Halo-ligand staining showing co-immunoprecipitation of GFP-tagged dNxt1 with Halo-tagged dNxf1 or dNxf2 from OSC cells. GFP serves as a negative control. **f,** Sequence alignment of the region of Panx that interacted with dNxf2. Residues involved in the interactions are marked by teal diamond.

### Extended Data Figure 5

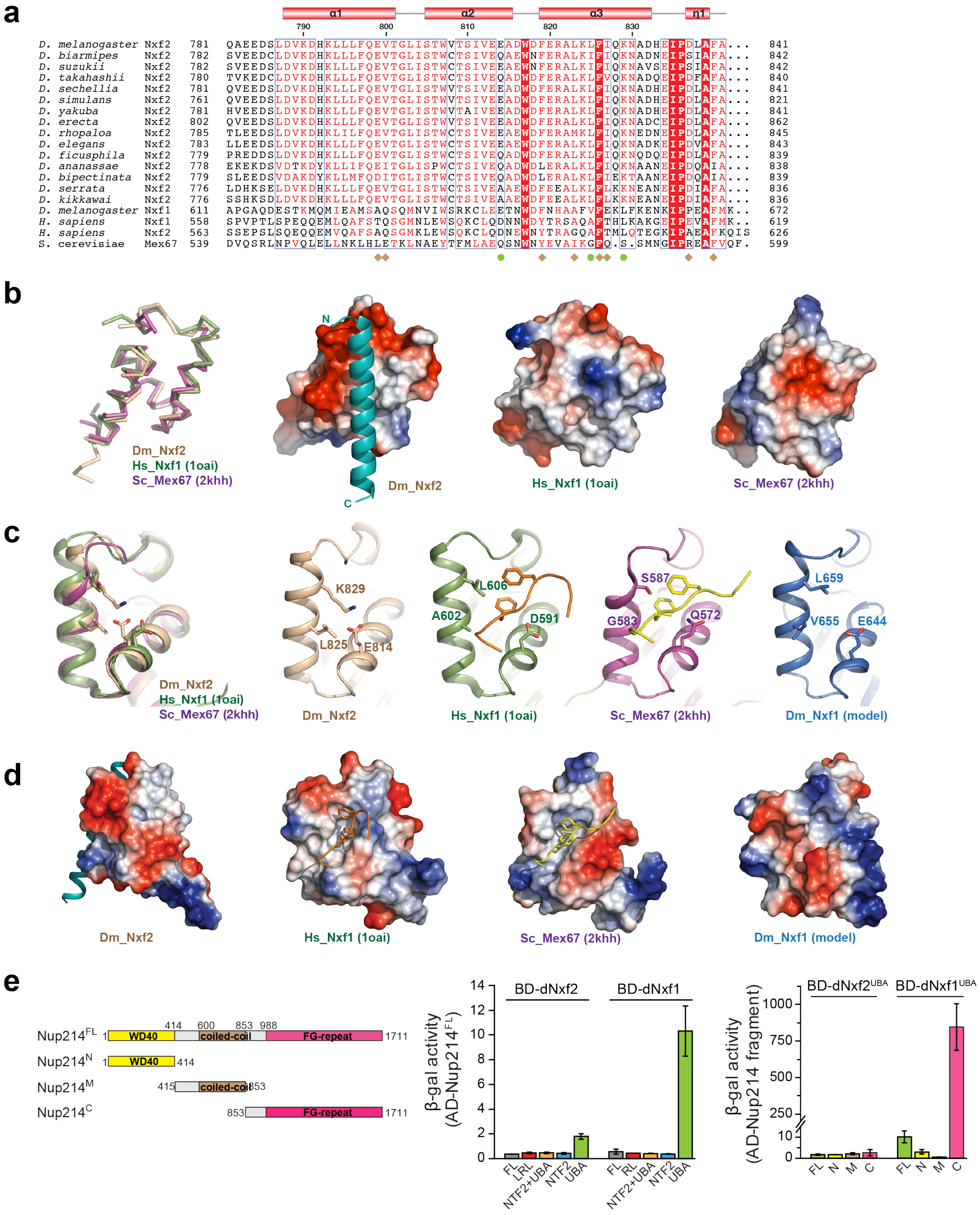

**Extended Data Figure 5. Structural comparison of the UBA domains from different NXF family proteins. a,** Sequence alignment of the UBA domains of NXF family proteins. Residues within the dNxf2-Panx interaction interface are indicated in brown diamond. Residues that blocks the entry of the FG peptides are indicated in green circle. Residues that are identical are shown in white with red shading. Residues with high similarities are shown in red. Secondary structural elements are indicated above. **b,** Superimposition of the UBA domains of dNxf2 (wheat), human Nxf1 (green, PDB ID: 1OAI), and yeast Mex67 (purple, PDB ID: 2KHH). Electrostatic potential analysis showing that the corresponding surfaces in the UBA domains of dNxf2 (left), human Nxf1 (middle) and yeast Mex67 (right) are positively charged and negatively charged, respectively. **Fig. 3c** was shown here again for comparison. **c,** Detailed view, left to right, of the UBA domains of dNxf2, human TAP, yeast Mex67, and dNxf1 (model). The dNxf1 model was generated by SWISS-MODEL server. In human Nxf1 and yeast Mex67, the FG peptides bind in the concave passage formed by α2 and α3. **d,** Electrostatic potential analysis showing a closed surface on dNxf2 UBA that blocks the entry of FG-nup. In the corresponding surface of dNxf2 UBA, the passage for the peptide is blocked (left). Human Nxf1 UBA (PDB ID: 1OAI) and yeast Mex67 UBA (PDB ID: 2KHH) show a hydrophobic pocket to accommodate the two phenylalanines of the FxFG peptides (middle). A model of dNxf1 UBA (right) indicates that a peptide binding pocket may exist on the dNxf1 surface. **e,** Y2H assays show that dNxf1 UBA domain but not dNxf2 UBA interacts with Nup214 FG-repeat region. Interactions were determined by either measuring the beta-galactosidase activity produced by the reporter gene. Data are averages of three independent experiments (n=3). Proteins or fragments of Nup214 are shown above the dashed line.

### Extended Data Figure 6

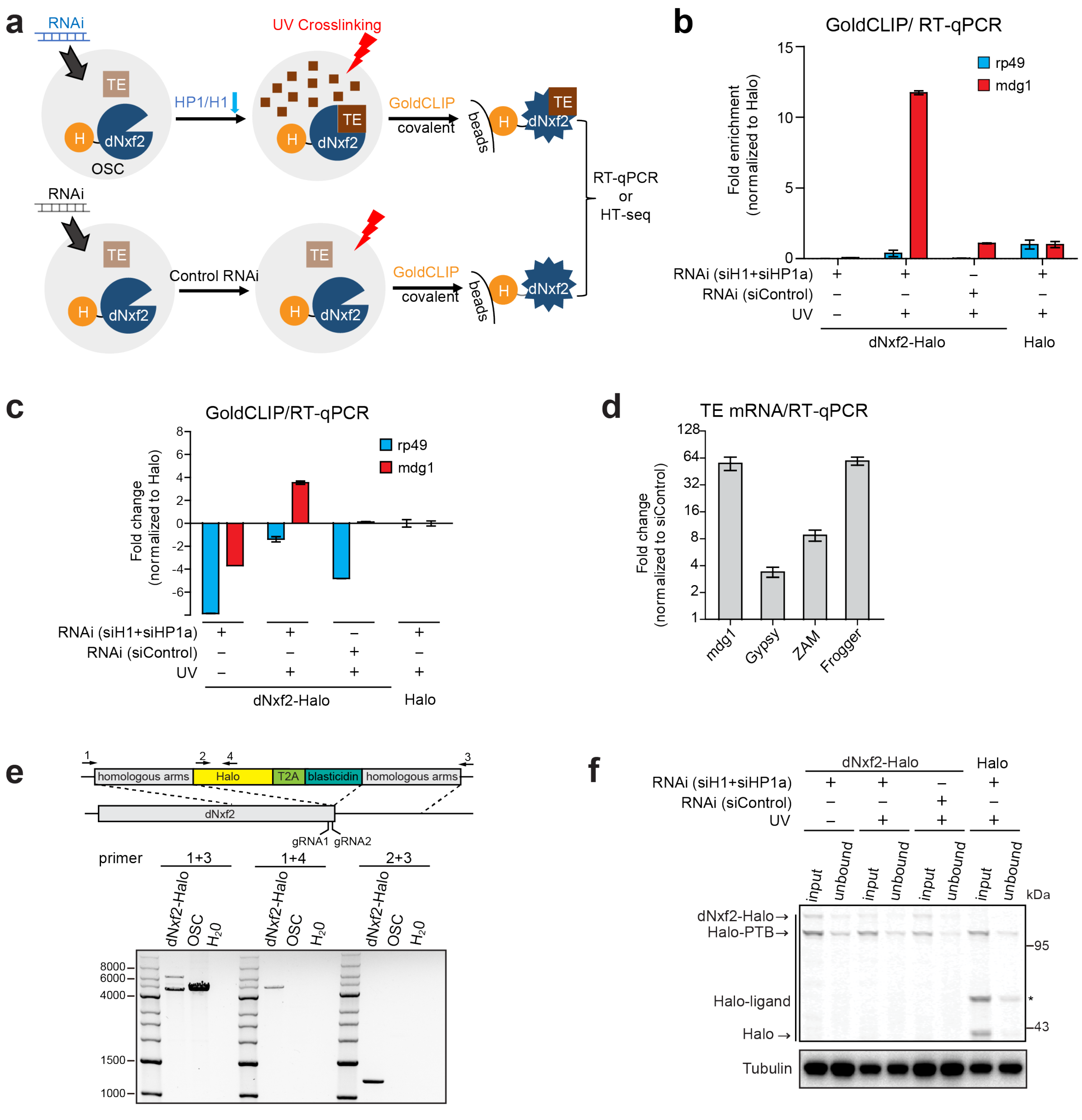


**Extended Data Figure 6. dNxf2 specifically binds to transposons *in vivo*. a,** Schematic representation of the GoldCLIP experiments. OSC, Ovarian Somatic Cell. TE, transposon. H, Halo-tag. **b,** RT-qPCR results showing relative levels of the mdg1 transposon or rp49 for dNxf2-Halo GoldCLIP sample, compared to either no UV or a Halo control. Fold enrichments are calculated as a mammalian spike-in normalized RNA levels divided by that of the input samples. For comparison, the value of Halo control was used for normalization. Mean values ± s.d. from 3 independent experiments are shown. **c,** Data from **b** was redrawn in $\log_2$ scale showing relative enrichment or depletion of the indicated transcripts, normalized to the Halo alone control. **d,** RT-qPCR results showing relative steady-state RNA levels of the indicated transposons for OSC from the H1+HP1a double siRNA knockdown, compared to a control siRNA knockdown. Fold changes are calculated as *rp49*-normalized RNA levels divided by that of the corresponding controls. Mean values ± s.d. from 3 independent experiments are shown. **e,** A Schematic of the knock-in strategy is displayed. Agarose gel showing PCR results confirming a correct integration of Halo-tag to the C-terminus of the dNxf2 locus. PCR primers amplifying the indicated regions are shown on the top and two sgRNAs targeting near the stop codon of dNxf2 are indicated at the bottom of the schematic. **f,** Western blots and Halo-ligand staining showing depletion of dNxf2-Halo or Halo alone by Halo pull down. Beta-tubulin blots serve as a loading control. The asterix indicates non-specific bands stained by Halo-ligand.

### Extended Data Figure 7

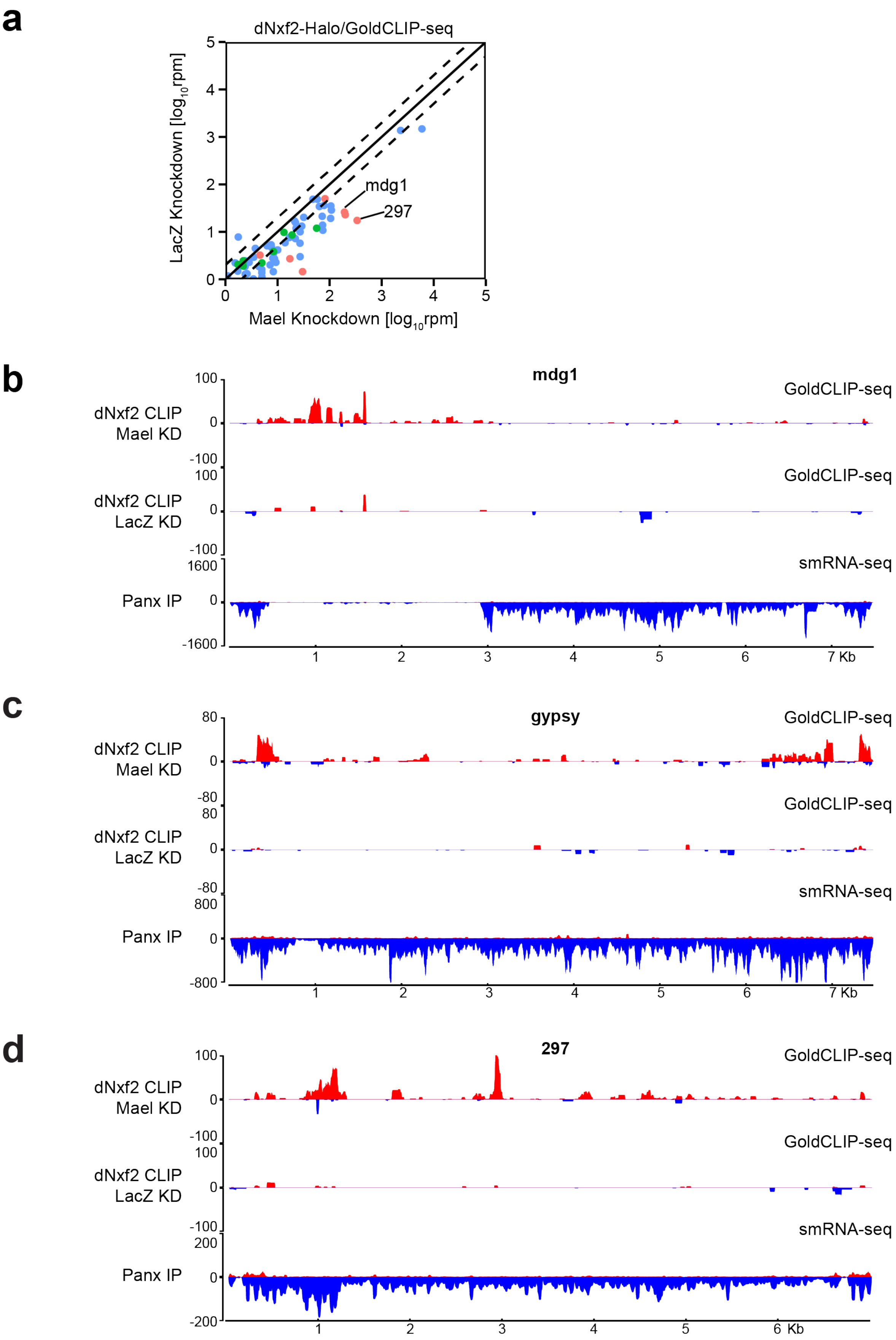


498

**Extended Data Figure 7. dNxf2 binds to Piwi/piRNA targeted transposons**. **a,** Comparison of GoldCLIP-seq are shown as reads per million (rpm) mapping to the sense strand of each transposon consensus for Maelstrom knockdown (*X* axis) versus a LacZ control knockdown (*Y* axis). A mammalian spike-in was used to normalize different samples. Dashed lines indicate two-fold changes. The pooled reads from two replicates are shown. Red dots indicate Piwi-targeted transposons, blue dots indicate H1-dependent transposons and green dots indicate other transposons. **b**, Density plots for dNxf2-Halo GoldCLIP reads from Mael knockdown OSCs (top panel), dNxf2-Halo GoldCLIP reads from LacZ knockdown OSCs (middle panel), and Panx co-immunoprecipitated smRNA reads (bottom panel, from previously published data[22]) over the transposon consensus of mdg1. **c** and **d** are the same as in **b** except for gypsy (**c**) and 297 (**d**).

### Extended Data Figure 8

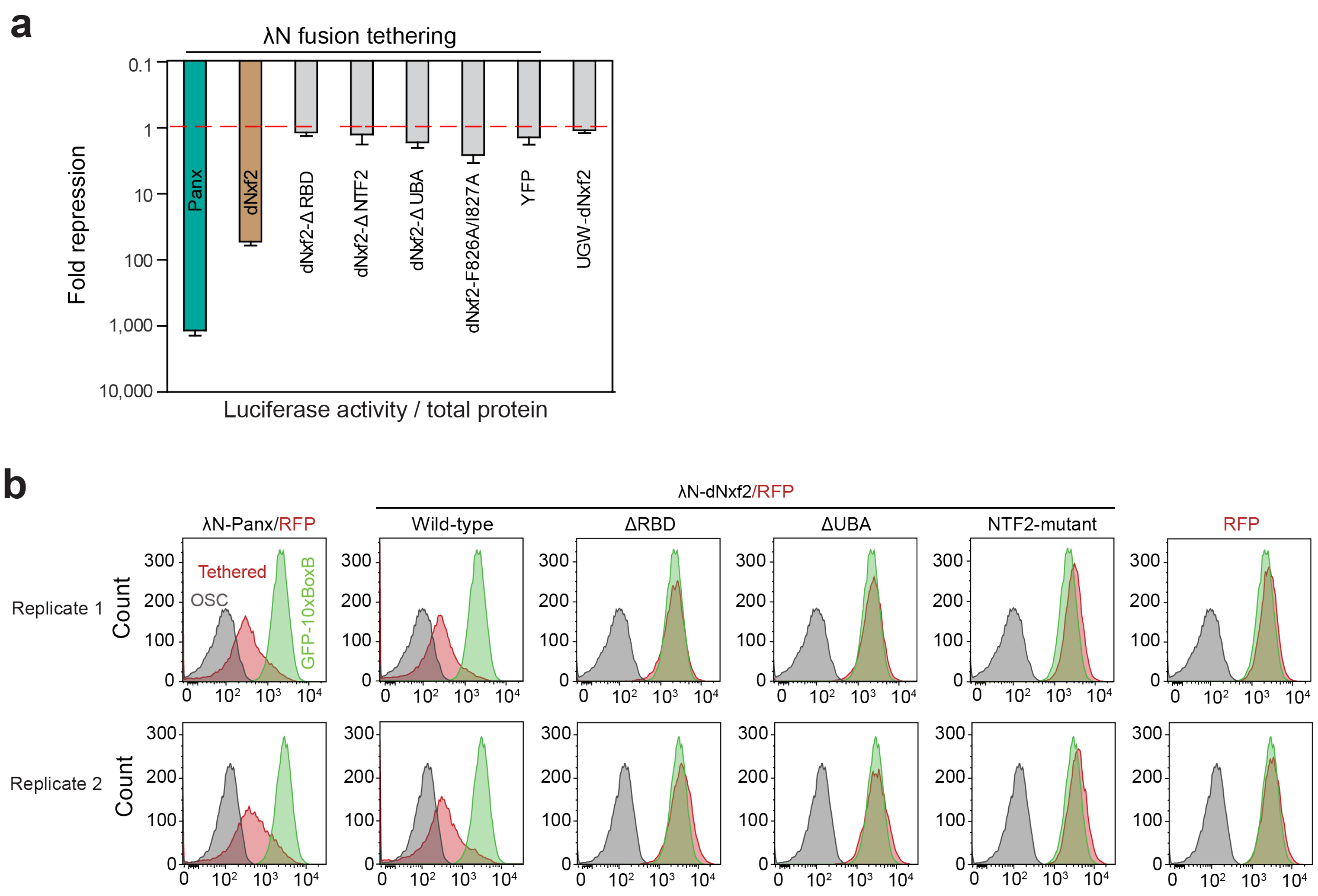


**Extended Data Figure 8. The Pandas complex is required for piRNA-guided transposon silencing. a,** Effects of the indicated λN fusion proteins or a non-tethering control (GFP-dNxf2) on luciferase activity of the reporters integrated into the attP2 landing site. Data from **Fig. 3g** was shown here again for comparison. Data show mean ± s.d. ($n$ = 15). **b,** FACS sorting showing counts and fluorescence of GFP positive cells gated by high RFP expressing OSCs electro-transfected with the indicated plasmids expressing different λN-fusion proteins. OSC (grey), autofluorescence of OSCs without any GFP reporter; Tethered (red), GFP reporter OSCs transfected with the indicated expression plasmids; GFP-10×BoxB (green), GFP reporter cells with no transfection.

### Extended Data Figure 9

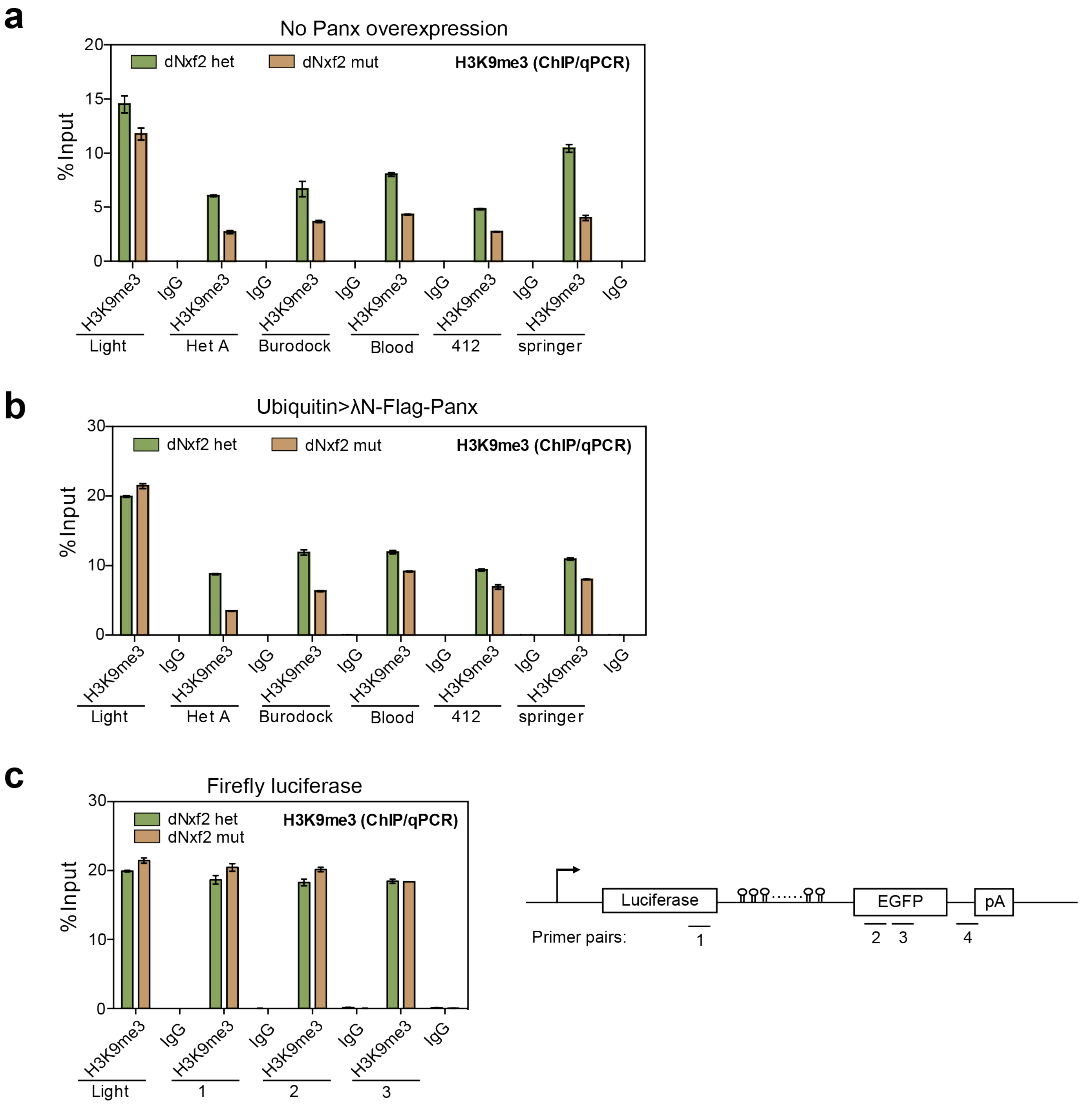


**Extended Data Figure 9. ChIP-qPCR assays measuring H3K9me3 changes upon loss of dNxf2. a,** H3K9me3 mark enrichments at transposons comparing heterozygous and mutant for dNxf2. **b,** H3K9me3 mark enrichments at the indicated transposons comparing heterozygous and mutant for dNxf2 when overexpressing Panx driven by a Ubiquitin promoter. Green and brown bar, polyclonal antibody specific to H3K9me3; Black and grey bar, pre-immune rabbit IgG. Mean values ± s.d. from 3 independent experiments are shown. **c,** H3K9me3 mark enrichments at the Firefly-10×BoxB reporter tethered with λN-flag-Panx in the same genetic background as in **b**. Schematic of qPCR primer pair positions is shown on the right.

### Extended Data Figure 10

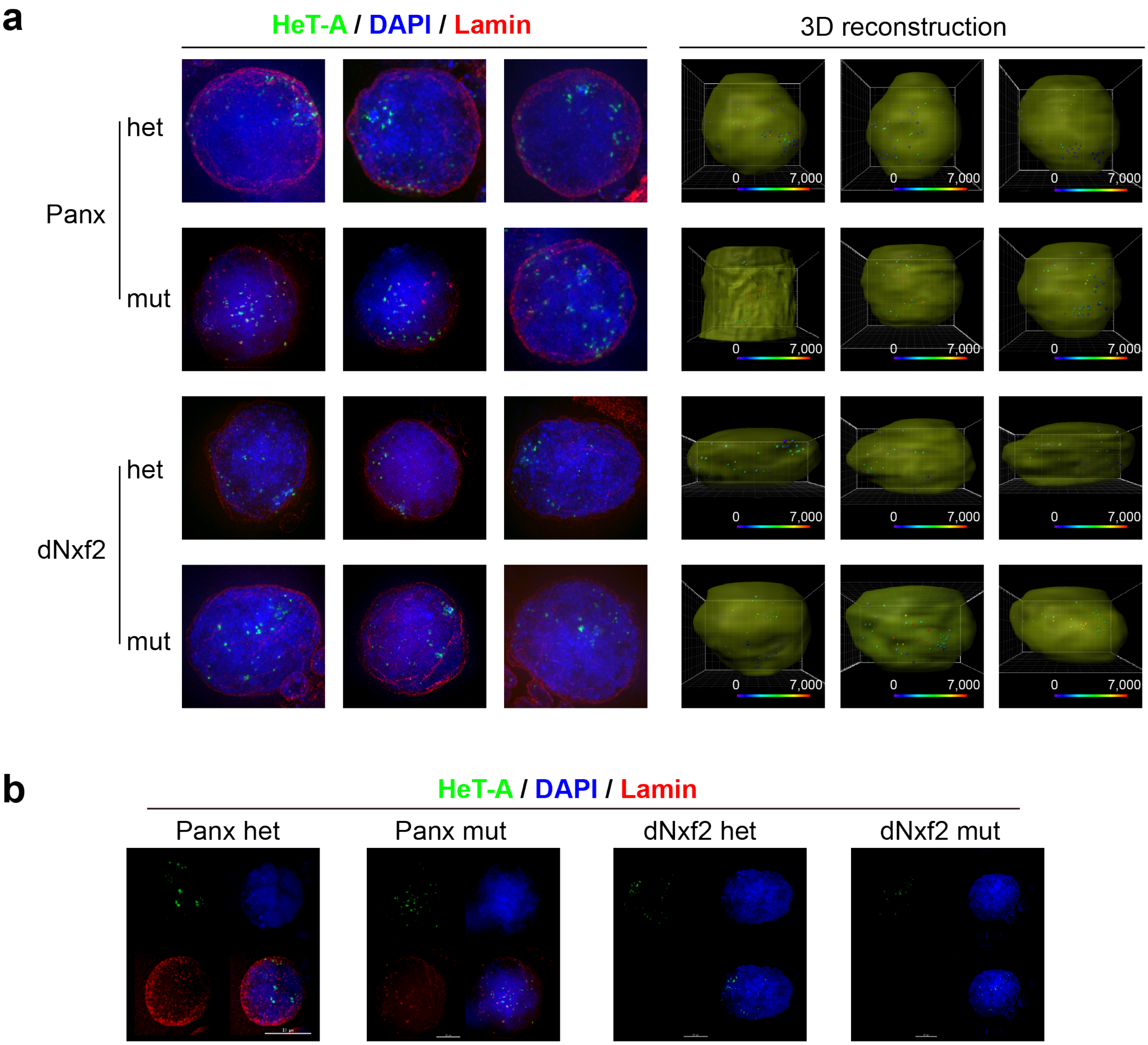


**Extended Data Figure 10. Het-A left nuclear peripheries upon loss of either Panx or dNxf2. a,** Left three columns showing 3D SIM super-resolution microscopy of DNA FISH (HeT-A, green) and Lamin A (red) double staining from either Panx or dNxf2 heterozygote versus mutant ovaries. Right three columns, estimation of the positioning of clustered HeT-A signals relative to the nuclear membrane of nurse cells. 10-20 nuclei were quantified for each condition. The distance was displayed in a blue-red scale with red indicating longer distance between the green dots and nuclear membrane. **b,** Projections of the 3D videos on the *XY* surface either in combination (bottom right) or in split channels for DAPI (blue, upper right), Lamin (red, bottom left) and HeT-A (green, upper left) on the selected ovaries from **a**. See also Supplementary Movies 1-4.

### Extended Data Figure 11

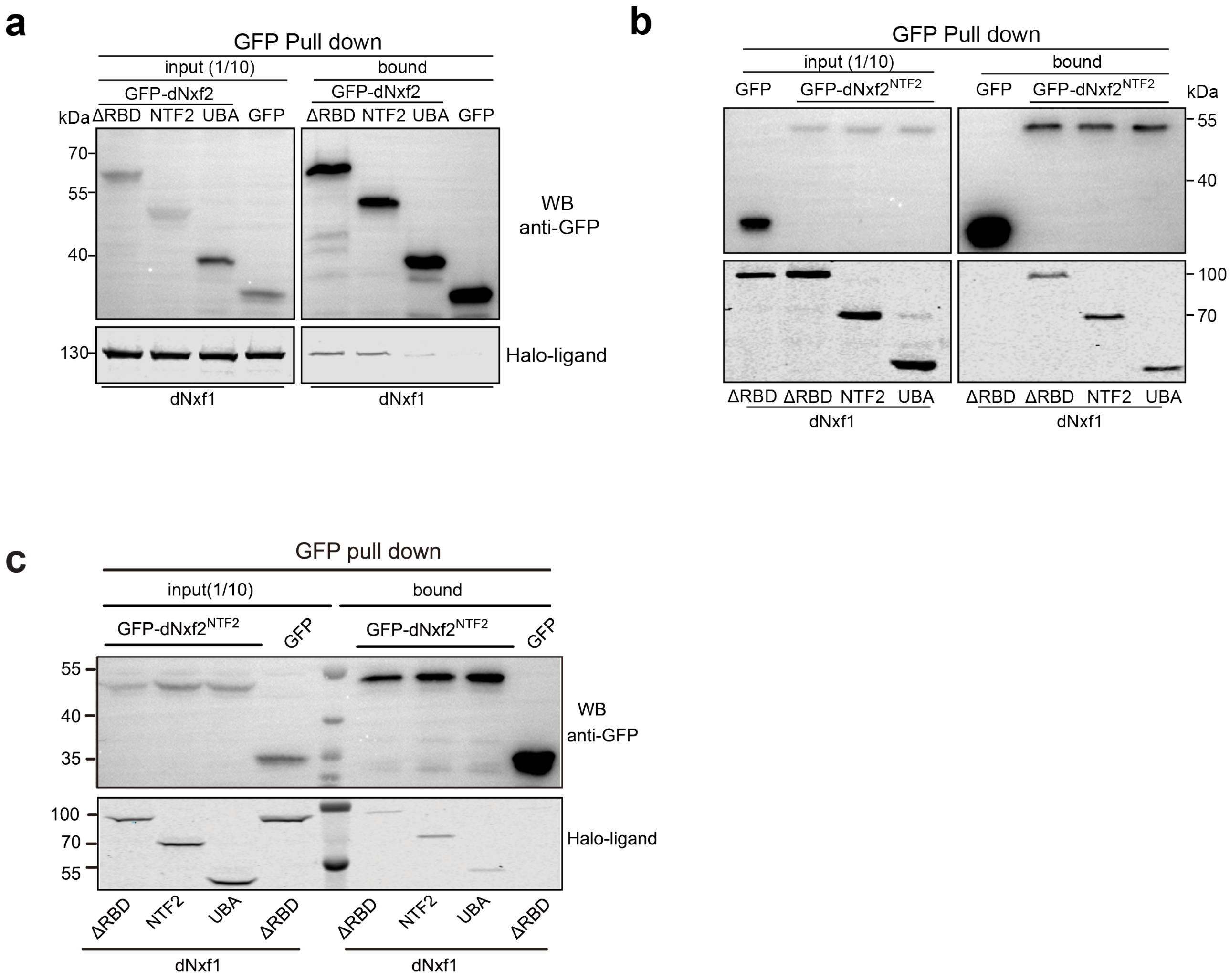


**Extended Data Figure 11. dNxf2 interacts with dNxf1 in *Drosophila* cell lines. a,** Western blots and Halo-ligand staining showing co-immunoprecipitation of GFP-tagged various truncated dNxf2 with Halo-tagged full length dNxf1 from OSC cells. ΔRBD, dNxf2 lacking N-terminus RNA binding domains; NTF2, dNxf2$^{NTF2}$ domain; UBA, dNxf2$^{UBA}$ domain. **b,** Western blots and Halo-ligand staining showing co-immunoprecipitation of GFP-tagged dNxf1$^{NTF2}$ domain with Halo-tagged various truncated dNxf1 from S2 cells. ΔRBD, dNxf1 lacking all N-terminus RNA binding domains; NTF2, dNxf1$^{NTF2}$ domain; UBA, dNxf1$^{UBA}$ domain. GFP serves as a negative control. **c,** The same as in **c** except the experiments were done using OSC cells.

### Extended Data Figure 12

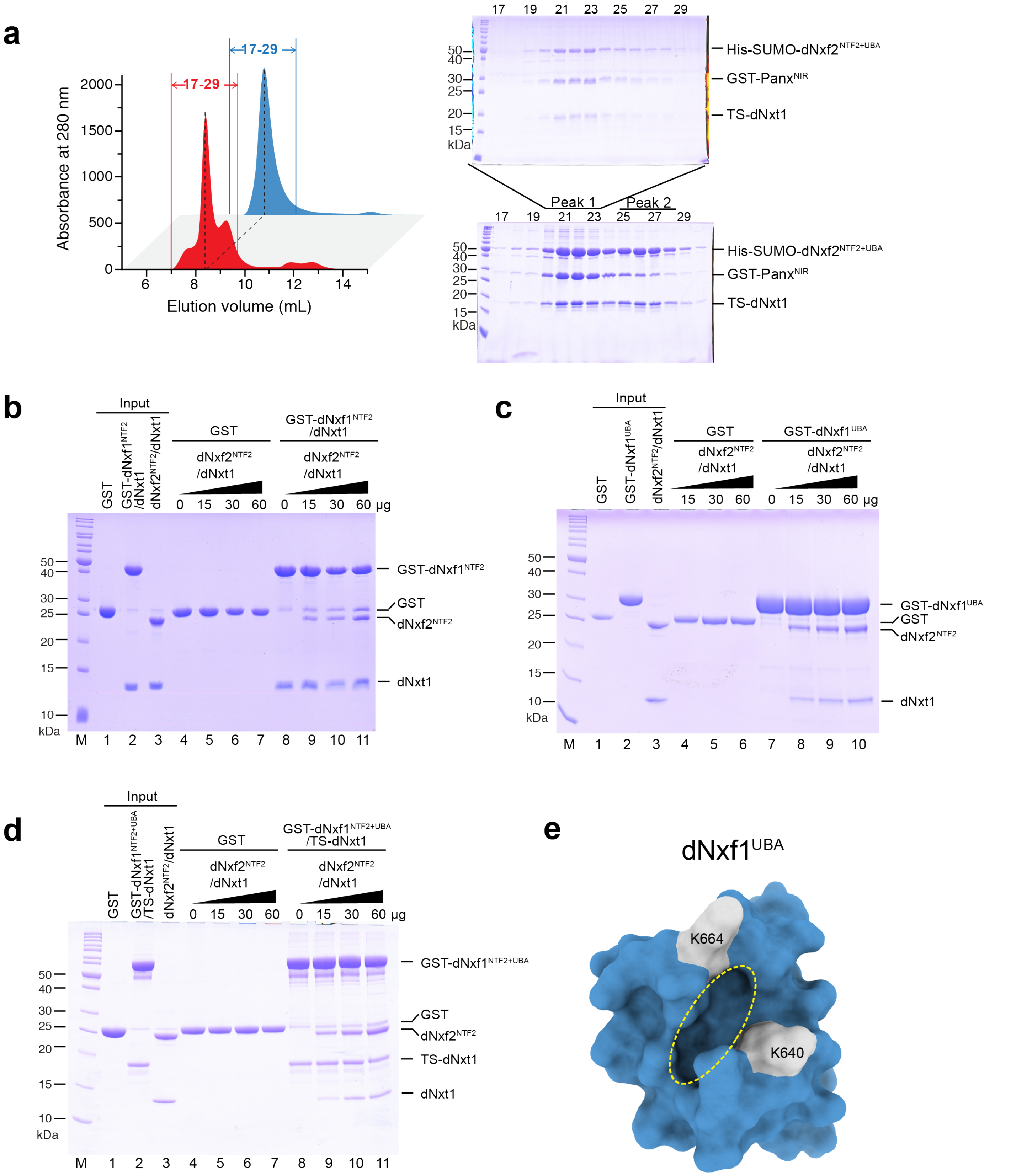


**Extended Data Figure 12. dNxf1 directly interacts with the NTF2 domain of dNxf2 via either NTF2 or UBA domains. a,** dNxf2$^{NTF2+UBA}$, dNxt1, and Panx$^{NIR}$ form a ternary complex in solution as shown on the size exclusion chromatography profile. Left panel, OD280 trace; Right panel, SDS-PAGE shows the components of the peak in the elution profile. **b-d,** GST pull-down assays showed the direct interactions between dNxf1 and dNxf2. Each of 60 μg of purified GST-dNxf1$^{NTF2}$/dNxt1 (**b**), GST-dNxf1$^{UBA}$ (**c**), GST-dNxf1$^{NTF2+UBA}$/TS-dNxt1 complex (**d**) and GST protein was immobilized on GSH beads, respectively. Increasing amounts of dNxf2$^{NTF2}$/dNxt1 complex (0, 15, 30, 60 μg) were added and incubated. The beads were washed and aliquots of the bound fractions (8%) were analyzed by SDS-PAGE. Each input protein of 2 μg was loaded on SDS-PAGE as well. Positions of molecular weight markers are indicated on the left in kDa. **e**, Schematic of dNxf2$^{NTF2}$-crosslinking residues K664 and K640 (white) on dNxf1$^{UBA}$ (blue). The dNxf1 model was generated by SWISS-MODEL server as shown in **Extended Data Fig. 5c**.The FG binding pocket is marked with a dash yellow circle.

### Extended Data Figure 13

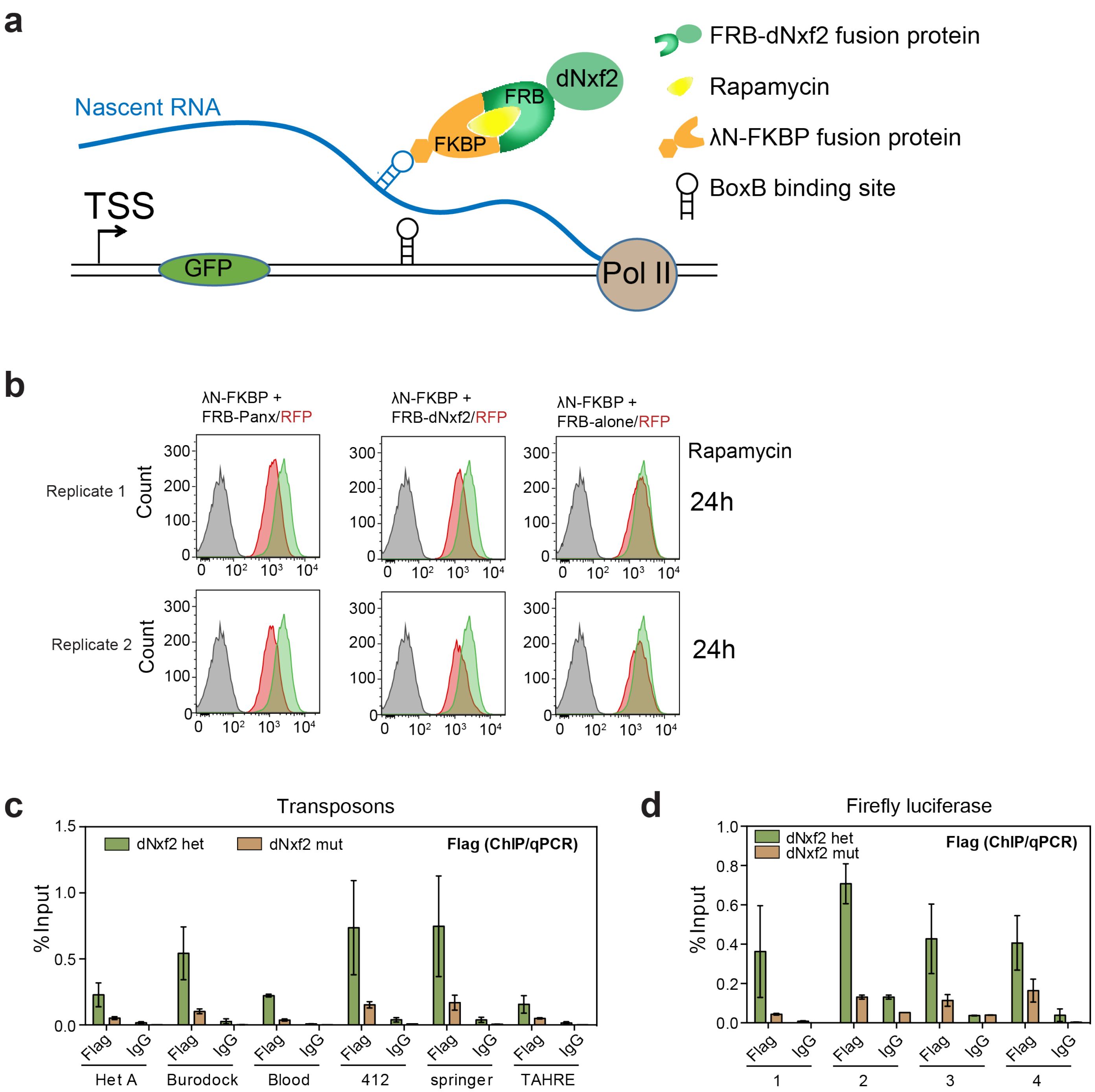


**Extended Data Figure 13. Rapamycin inducible tethering system. a,** Schematic representation of a rapamycin-inducible system for λN-FKBP:FRB tethering. **b,** FACS sorting showing counts and fluorescence of GFP positive cells gated by high RFP expressing OSCs (as in **Extended Data Fig. 7b**) transfected with the plasmids expressing both λN-FKBP and the indicated FRB-fusion proteins. 24 hrs after transfection, cells were treated with rapamycin for additional 24 hrs. Two independent biological replicates were shown. **c,** Flag-Panx enrichments at the indicated transposons comparing heterozygous and mutant for dNxf2 when overexpressing Panx driven by a Ubiquitin promoter. Green and brown bar, monoclone antibody specific to Flag (M2); Black and grey bar, pre-immune mouse IgG. Mean values ± s.d. from 3 independent experiments are shown. **d,** Flag-Panx enrichments at the Firefly-10×BoxB reporter tethered with λN-flag-Panx in the same genetic background as in (**c**). Schematic of qPCR primer pair positions is shown in **Extended Data Fig. 9c**.
